# BEST4/CA7^+^ and goblet cells are interdependent regulators of intestinal mucus homeostasis

**DOI:** 10.1101/2025.09.29.679286

**Authors:** Willem Kasper Spoelstra, Daisong Wang, Johan H. van Es, Hans Clevers, Jeroen S. van Zon, Sander J. Tans

## Abstract

The intestinal mucus layer is essential for the integrity of the intestinal barrier. It is produced by goblet cells, whose depletion is common in colonic inflammation but remains poorly understood. Here, we show that goblet cell survival relies on a reciprocal dependence with newly discovered BEST4/CA7^+^ cells. We developed a method to follow BEST4/CA7^+^ and goblet cells in time from birth to death in human colon organoids. Notably, goblet cells induce BEST4/CA7^+^ fates in sister cells and other neighbors, using DLL1-mediated lateral activation of Notch-signaling. BEST4/CA7^+^ cells in turn promote goblet survival, with the latter depleting rapidly after differentiation in absence of BEST4/CA7^+^ cells. This apoptosis inhibition does not require direct cell-cell contact and instead depends on their shared lumen. Such differentiation and survival interdependencies may be relevant beyond the maintenance of mucosal homeostasis.

---

The intestinal lumen is a hostile environment that contains diverse pathogens, which must be effectively separated from the delicate underlying tissue. A mucus layer covering the intestinal epithelium is central to this barrier function. It consists mostly of the *O*-glycosylated mucin 2 (MUC2), which expands up to 1000-fold to form a hydrated gel after its secretion by goblet cells^1–3^. Emerging evidence also indicate a role for BEST4/CA7^+^ cells; a rare cell type discovered only recently in the human intestine^4^ and notably absent in the mouse^5^. Specifically, the key markers BEST4 and CA7 both regulate bicarbonate levels. BEST4/CA7^+^ cells also express proton channels OTOP2 and OTOP3^6–11^, secrete guanylin and uroguanylin^9^, and activate CFTR channels upon bacterial infection^12^. These findings indicate that BEST4/CA7^+^ cells regulate mucus viscoelasticity and secretion by controlling luminal pH, calcium and bicarbonate concentrations, as well as water efflux^13–15^ (see ref. ^16^ for recent review).

Goblet and BEST4/CA7^+^ cell abundance is therefore important to mucosal defense. Indeed, goblet cell depletion is a known hallmark of ulcerative colitis^17^, and more generally observed in chronic inflammatory disorders^18–21^. However, the processes that control goblet and BEST4/CA7^+^ cell abundance in the rapidly renewing intestinal tissue are poorly understood. Goblet cells arise from secretory progenitors, but whether their survival is controlled is less clear. BEST4/CA7^+^ cells are thought to branch from absorptive progenitors^9^ and are absent in organoid cultures treated with Notch signaling inhibitors^12,22^. However, when and in which lineages they are generated is unknown. Curiously, BEST4/CA7^+^ and goblet cells are often found in close proximity^4,6^, but the causes are unclear. Addressing these questions requires approaches to follow the renewal process of intestinal cells in time and space. Owing to the absence of BEST4/CA7^+^ cells in the mouse, studying their differentiation and function over time requires the use of human organoid models^12,22^.

Here, we developed a method to track goblet and BEST4/CA7^+^ cells in a human colon organoid model. The data reveal an unexpected interdependence between these cell types in terms of both differentiation and survival. Notably, we find that BEST4/CA7^+^ cells emerge often, but not exclusively, as direct sisters of goblet cells. Fate commitment occurs early, with goblet cells arising from cells that are still proliferative, and their sister or other neighboring cells adopting the BEST4/CA7^+^ fate shortly after. We find that goblet cells laterally activate Notch-signaling through DLL1 to induce BEST4/CA7^+^ cell differentiation. Finally, we show that goblet cell survival depends on BEST4/CA7^+^ cells in a contact-independent manner, with the goblet cell pool being dramatically depleted in absence of BEST4/CA7^+^ cells. This reciprocal relation between goblet and BEST4/CA7^+^ cells provides a mechanism for homeostasis of the ratio between BEST4/CA7^+^ and goblet cells. Our findings raise questions on the prevalence of reciprocal dependencies between cell types and their role in regulating cell type proportions across epithelia and during infection, inflammation, and proliferative perturbations.

## Results

### BEST4/CA7^+^ cells differentiate after goblet cells

To study the dynamics of goblet and BEST4/CA7^+^ cell differentiation, we performed long-term live-cell imaging of human colon organoids^23^ carrying reporters for differentiation of goblet cells (labelled by a *MUC2-mNeonGreen* knock-in allele)^24^ and BEST4/CA7^+^ cells (labelled by a *CA7-P2A-tdTomato* knock-in allele)^12^, with a histone 2B (H2B)-iRFP670 nuclear reporter integrated to enable real-time tracking of individual cells. We grew human colon organoids as undifferentiated stem cells in expansion medium and then shifted to differentiation medium (**Fig. 1a**). Multi-day time-lapses were acquired during these growth processes at 12-min resolution using confocal microscopy and individual nuclei were tracked through time using OrganoidTracker^25^. We classified cells as goblet cells, BEST4/CA7^+^ cells or unmarked cells based on morphology and fluorescence, and extracted cell positions, fluorescence levels of mNeonGreen and tdTomato (which indicate expression levels of MUC2 and CA7, respectively) and lineage relations (**Online Methods, Fig. 1a-d and Extended Data** Fig. 1,2). Together, this yielded an expansive single cell tracking dataset combining positional information, lineage history, gene expression dynamics and cell types.

**Fig. 1:**
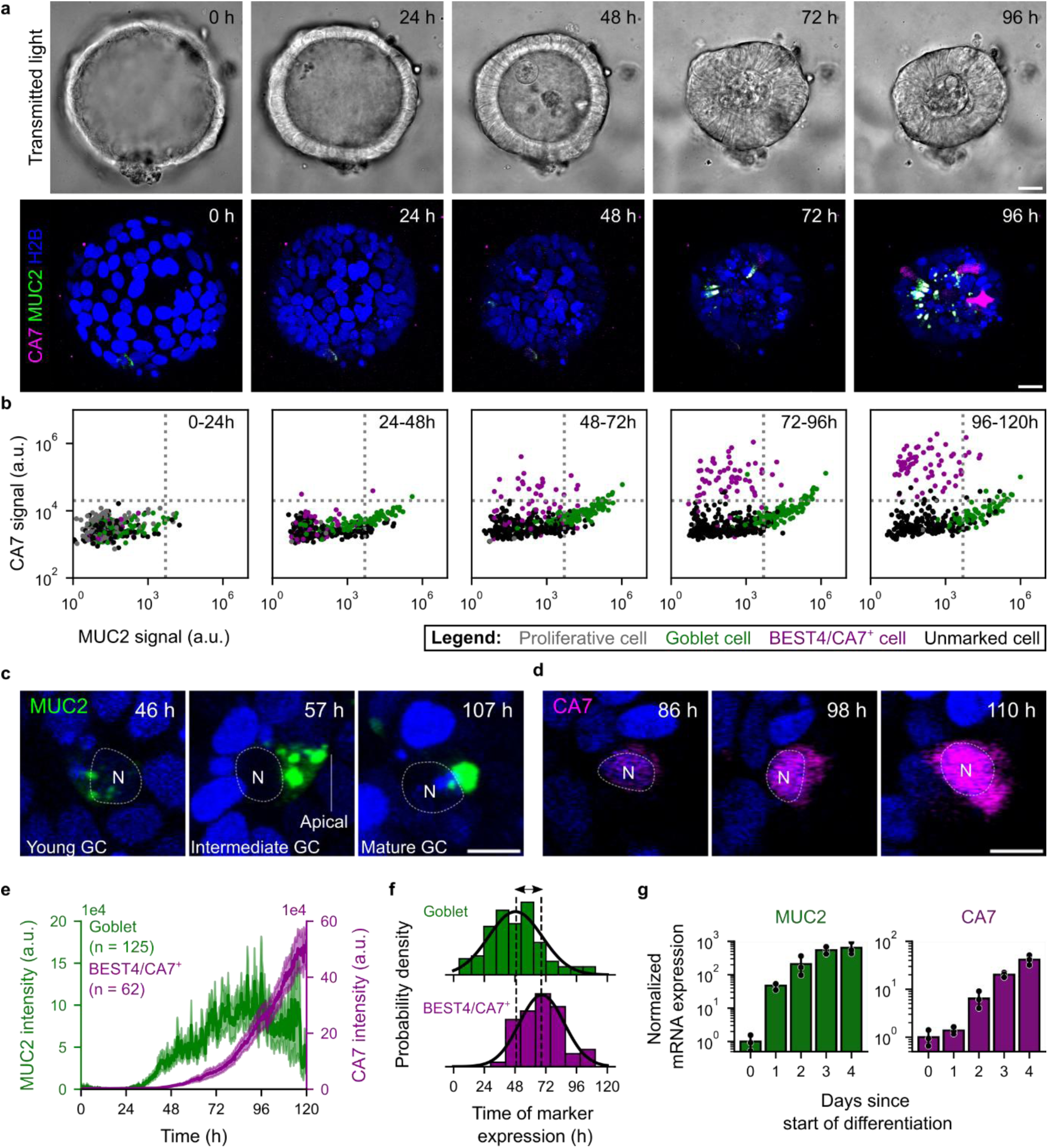
Goblet cells emerge before BEST4/CA7^+^ cells. a, Transmitted light (top) and fluorescence micrographs of a differentiating CA7-MUC2-H2B triple-reporter organoid. Time is hours after shift to differentiation medium. Scale bars: 20 µm. **b,** MUC2 (x-axis) and CA7 (y-axis) reporter fluorescence averaged over subsequent time windows. **c,d,** Maturation of a goblet cell (GC, **c)** and BEST4/CA7^+^ cell **(d)**. Scale bars: 10 µm. **e,** Average of MUC2 intensity for all goblet cells (green, left y-axis) and average CA7 expression of BEST4/CA7^+^ cells (magenta, right y-axis). 17 organoids derived from three independent experiments are analyzed. n indicates the number of analyzed cells; shaded area indicates the S.E.M. **f,** Distribution of times at which *MUC2* (top) and *CA7* (bottom) is first expressed. Vertical dashed lines indicate the median expression times. Solid line indicates a normal distribution fit to the histogram. **g,** Expression of *MUC2* (green, left) and *CA7* (magenta, right) mRNA quantified using qPCR, at different days after shifting to differentiation medium. Vertical axis indicates the expression relative to *GAPDH*, normalized by the average expression at day 0 (data from three biological replicates).

Lineage trees showed that cells in expansion medium remained proliferative for at least five days with cell cycle durations varying between 15 and 53 hours, and a median of 27.6 hours (**Extended Data** Fig. 1a-e **and Supplementary Video 1**). Cell cycle durations were strongly correlated between sister cells (**Extended Data** Fig. 1f), in line with our previous finding in mouse intestinal organoids that cell cycle durations are controlled by mother cells and transmitted to their daughters^26^. After shifting to differentiation medium, most cell divisions occurred within 24 hours and proliferation virtually stopped within 48 hours (**Extended Data** Fig. 1g). After 24 hours, a subset of cells started to show *MUC2* and *CA7* expression (**Fig. 1e and Supplementary Video 2**). Notably, we observed a distinct temporal order between goblet and BEST4/CA7^+^ cell emergence, with MUC2^+^ goblet cells appearing after a median time of 49 hours, and BEST4/CA7^+^ cells after 69 hours (**Fig. 1e,f**). Measurements of *MUC2* and *CA7* mRNA levels by quantitative polymerase chain reaction (qPCR) analysis yielded a similar delay in timing (**Fig. 1g**). This temporal separation can explain previous immunostaining^27^ and transcriptomics data^7,9^, which show BEST4/CA7^+^ cells higher along the crypt/crypt-villus axis than goblet cells, as the former will have moved further towards the villus before their BEST4/CA7 marker is expressed.

### Contact with goblet cells predicts BEST4/CA7^+^ cell fate commitment

BEST4/CA7^+^ cells often appear to be positioned nearby goblet cells both *in vivo* and *in vitro*^4,6,12^. Rigorously identifying cell-cell contact in 3D tissue architectures is challenging, however. To address this issue quantitatively, we grew organoids in 2D monolayer cultures^28^, to ensure that cell interfaces are clearly visible in bright-field images (**Fig. 2a,b**). On day 3 of differentiation, 90% of BEST4/CA7^+^ cells were in direct contact with goblet cells. In contrast, only 44% of goblet cells and 42% of unmarked cells were in contact with another goblet cell (**Fig. 2c,d**). Furthermore, sister cells remained close together after division in both expansion and differentiation medium (**Extended Data** Fig. 3). These findings excluded the possibility that goblet and BEST4/CA7^+^ cells differentiate independently and subsequently co-localize by spatial rearrangement. Instead, lineage relations or cell-cell signaling could play a role.

**Fig. 2:**
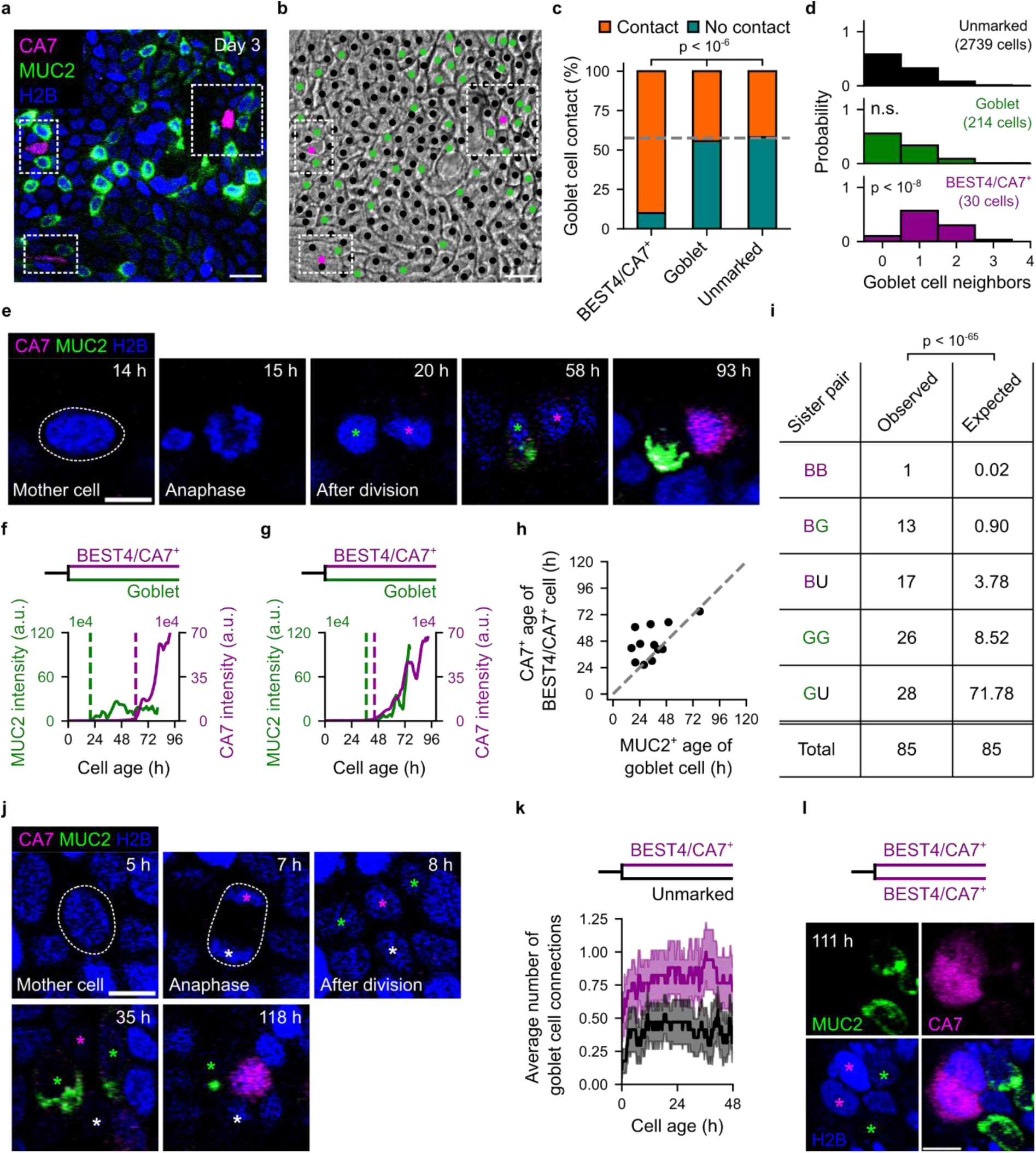
Contact with goblet cells predicts BEST4/CA7^+^ cell fate. a,b, Fluorescence **(a)** and bright-field **(b)** micrograph showing goblet-BEST4/CA7^+^ cell co-localization in 2D organoids. Dashed white boxes highlight BEST4/CA7^+^ cells. Scale bars: 20 µm. **c,** Fraction of cells in contact with a goblet cell. Pearson χ^2^-test is used for comparisons. n = 17 patches from three independent experiments. **d,** Distribution of the number of goblet cell neighbors per cell type. Mann-Whitney U test is used for comparisons. n.s. not significant. n = 17 patches from three independent experiments. **e,** Example of a dividing cell of which one daughter becomes a goblet cell (green asterisk) and the other a BEST4/CA7^+^ cell (magenta asterisk). Time is hours after shift to differentiation medium. Scale bar, 10 µm. **f,g,** Two representative examples for *MUC2* expression of goblet cells (green) and *CA7* expression of their BEST4/CA7^+^ sister. Age indicates time since the last division. Vertical dashed lines indicate the time at which the cells became MUC2-positive (green) or CA7-positive (magenta). Both lines show the 4-hour moving average of the MUC2-mNeonGreen and CA7-P2A-tdTomato fluorescence. **h,** Age at which goblet and BEST4/CA7^+^ cells showed detectable levels of marker expression in goblet- BEST4/CA7^+^ sister pairs. **i,** Observed and expected frequencies of the sister pairs. Expected frequencies were computed using the population-level abundances of goblet and BEST4/CA7^+^ cells and tested using a Pearson χ^2^-test. B, BEST4/CA7^+^ cell; G, goblet cell; U, unmarked cell. **j,** Example of a mother cell dividing into a BEST4/CA7^+^ and unmarked cell, indicated by a magenta and white asterisk, respectively. Green asterisks indicate MUC2^+^ cells adjacent to the BEST4/CA7^+^ cell. Scale bar: 10 µm. **k,** Average number of goblet cell connections for BEST4/CA7^+^ cells and unmarked BEST4/CA7^+^ cell sisters. A goblet cell connection is defined as an instance where the nucleus of a cell is within 15 μm of the nucleus of a goblet cell. Shaded area indicates the S.E.M. **l,** Micrograph images of the only BEST4/CA7^+^-BEST4/CA7^+^ sister pair, showing that both BEST4/CA7^+^ sisters (magenta asterisks) are in contact with a goblet cell (green asterisks). Scale bar: 10 µm.

To study this issue directly, we analyzed goblet and BEST4/CA7^+^ lineages in time, as well as their relations. We excluded lineages with cells whose type could not be classified as BEST4/CA7^+^, goblet, or not expressing either marker (termed unmarked), for example due to cell death or the cell moving out of view (**Supplementary Note 1 and Extended Data** Fig. 4). Strikingly, we observed cells dividing into two sister cells of which one adopted the goblet, and the other the BEST4/CA7^+^ cell fate (**Fig. 2e**). Expression of the goblet cell marker *MUC2* typically turned on before the BEST4/CA7^+^ marker *CA7*, and never after (**Fig. 2e-h**), consistent with the population-level findings (**Fig. 1e-g**). Hence, these data show that goblet and BEST4/CA7^+^ lineages can emerge from the same progenitor (mother) cell. Overall, we observed sister cells displaying all five possible combinations that include BEST4/CA7^+^ and/or goblet type (**Fig. 2i and Extended Data** Fig. 2c). Analysis showed that the goblet- BEST4/CA7^+^ combination appeared more often than expected for a model where cells choose their fate in an autonomous and neighbor-independent manner (**Fig. 2i and Supplementary Note 2**).

The data further revealed a key role for spatial relations. Specifically, we considered cases when BEST4/CA7^+^ cells did not emerge as a goblet cell sister, but rather as a sister of an unmarked cell. We previously established that almost all unmarked cells in these organoids are enterocytes^12^, which are also dominant among differentiated cells in the colon. In unmarked- BEST4/CA7^+^ sister pairs, the BEST4/CA7^+^ cell typically had more contact with goblet cells than its unmarked sister cell (**Fig. 2j,k**). In the only observed sister pair with two BEST4/CA7^+^ cells, both BEST4/CA7^+^ sisters contacted goblet cells continuously until they expressed CA7 (**Fig. 2l**). Together, these results suggested a mechanism in which direct goblet cell contact instructs the BEST4/CA7^+^ cell fate. This model is consistent with all the above observations: the high frequency of goblet- BEST4/CA7^+^ sister pairs which contact each other after birth, the temporal order of their differentiation markers, and the induction of the BEST4/CA7^+^ fate in cells that contact goblet cells of a different lineage.

The data indicated another notable feature, namely the emergence of the goblet fate in each of the two sisters after a division. Indeed, goblet-goblet sister pairs were ∼3-fold overrepresented compared to expectation (**Fig. 2i**). In addition, the onset of *MUC2* expression relative to the last division was highly correlated for these goblet cell sisters (**Extended Data** Fig. 5a), suggesting that the mother cell had already committed to the goblet fate and then divided once more to produce two daughters in which MUC2 became detectable. Indeed, we occasionally observed the onset of *MUC2* expression already before this last division (**Extended Data** Fig. 5b). These observations are consistent with our previous study in mouse small intestinal organoids that showed secretory cells emerging as sisters^29^ with the same type, and oppose the view that secretory fate commitment occurs exclusively after cell cycle exit.

### BEST4/CA7^+^ cell fate commitment depends on DLL1 from goblet cells

The data thus far suggested goblet cells instruct the BEST4/CA7^+^ fate in direct neighbors. We surmised it could be mediated by Notch-signaling; a contact-dependent signaling pathway that regulates fate specification in the intestinal epithelium^30,31^. RNA-sequencing has shown that BEST4/CA7^+^ cells highly express the NOTCH2 receptor^6,7,9,10,12^, while inhibition of Notch-signaling using the γ-secretase inhibitor DAPT blocked all differentiation of BEST4/CA7^+^ cells^12,22^. However, as DAPT uniformly blocks γ-secretase activity along all lineages, the role of Notch signaling in BEST4/CA7^+^ specification, including the involvement of goblet cells, remains unclear. Therefore, we aimed to test directly whether goblet cells instruct the BEST4/CA7^+^ cell fate via lateral Notch-signaling.

In human tissue, five Notch-ligands are expressed, namely Delta-like ligands DLL1, DLL3 and DLL4 and Serrate-like ligands JAG1 and JAG2^32^. To identify candidate Notch-ligands for activation of Notch signaling in BEST4/CA7^+^ cells, we analyzed single-cell RNA sequencing data of healthy human colon epithelium from the GutCellAtlas^6^. We found that DLL3 and JAG2 were lowly expressed and DLL1 and DLL4 were highly expressed by colonic goblet cells (**Extended Data** Fig. 6). JAG1 was previously found to be expressed only in crypt top colonocytes^33^, and was therefore unlikely to contribute to Notch signaling in the differentiation process of BEST4/CA7^+^ cells. Although DLL4 shows overall higher expression than DLL1 (**Extended Data** Fig. 6), the latter was previously found to have higher affinity for NOTCH2 than DLL4^34^, suggesting distinct but not redundant role of Notch ligands in signaling activation. Hence, we hypothesized that any Notch signaling that instructs BEST4/CA7^+^ cell fate commitment is activated by DLL1, DLL4 or both.

To test this, we generated organoid lines where either *DLL1*, *DLL4* or both *DLL1* and *DLL4* were genetically knocked out on both alleles in the *CA7-P2A-tdTomato* reporter organoids (*DLL1*-KO, *DLL4*-KO and *DLL1*/*4*-dKO, respectively; **Extended Data** Fig. 7a). After organoid differentiation, we measured the percentage of BEST4/CA7^+^ cells by FACS analysis (**Extended Data** Fig. 7b). *DLL1*-KO and *DLL1*/*4*-dKO organoids had virtually no BEST4/CA7^+^ cells, while the percentage of BEST4/CA7^+^ cells was only mildly decreased in *DLL4*-KO organoids (**Fig. 3a,b**). Hence, expression of *DLL1*, but not *DLL4*, is necessary for instructing BEST4/CA7^+^ cell fate.

**Fig. 3:**
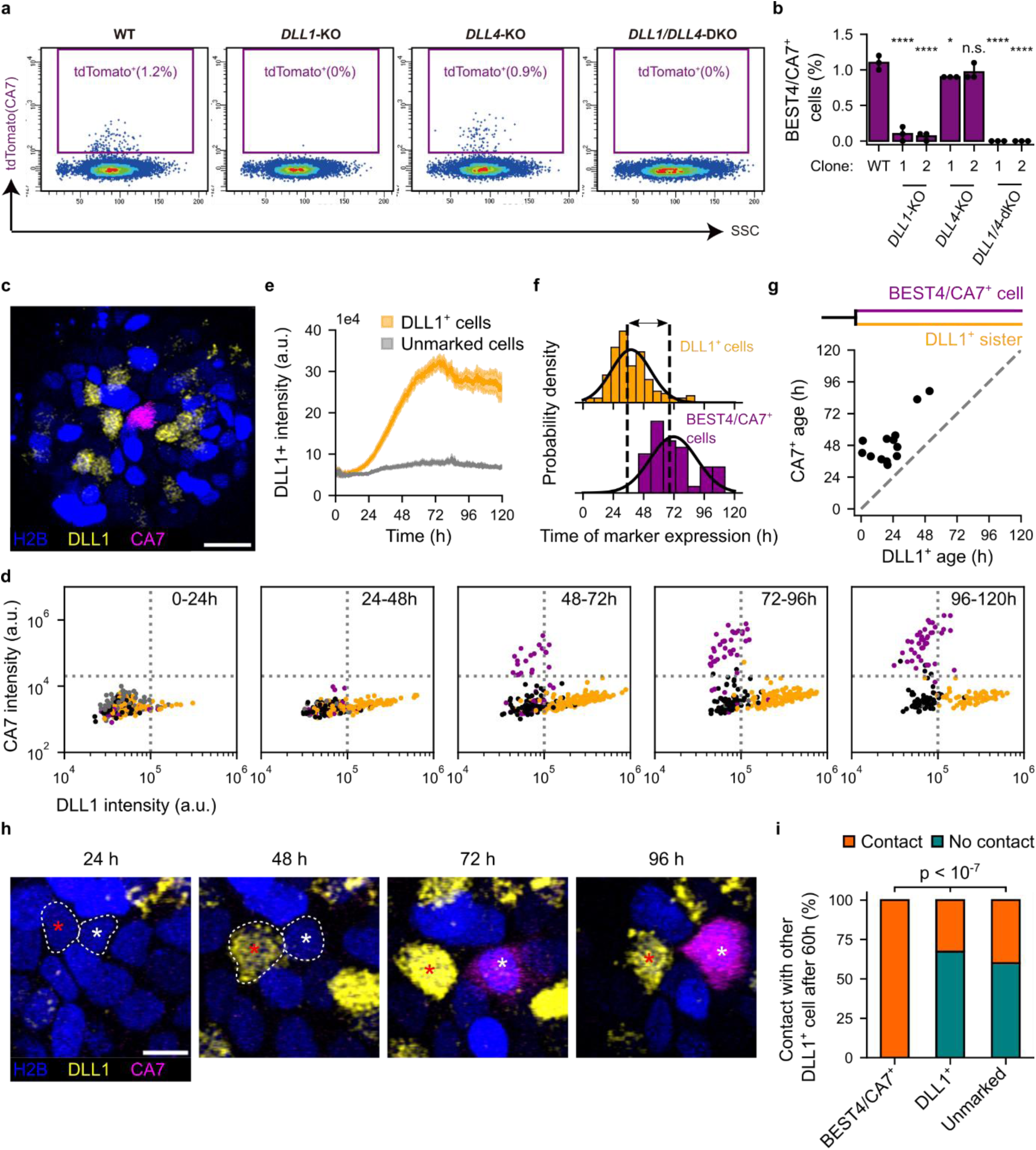
DLL1-mediated activation of Notch-signaling instructs BEST4/CA7^+^ cell fate. a,b, Representative FACS plot **(a)** and quantification of BEST4/CA7^+^ cell percentages **(b)** in differentiated WT, *DLL1*-KO, *DLL4*-KO and *DLL1*/*4*-dKO organoids. One-way ANOVA with Dunnett’s test is used for multiple comparisons between the DLL-KO groups and the WT (control) group. **** p < 0.0001, * p < 0.05, n.s.: not significant. SSC, side scatter measurement. Data from n = 3 technical replicates per condition, from N = 2 clones per knock- out line derived from the same donor. Error bars indicate 95% confidence interval. **c,** Representative confocal image of a DLL1-CA7-H2B reporter organoid, after 84 hours of differentiation. Scale bar: 20 µm. **d,** DLL1 (x-axis) and CA7 (y-axis) reporter fluorescence averaged over time for different time windows after shifting to differentiation medium. **e,** DLL1 reporter fluorescence over time for DLL1^+^ cells and unmarked cells. Shaded area indicates S.E.M. **f,** Distribution of times at which *DLL1* (top) and *CA7* (bottom) is first expressed. Vertical dashed lines indicate the median expression times. Solid line indicates a normal distribution fit to the histogram. **g,** Age at which BEST4/CA7^+^ and DLL1^+^ cells in BEST4/CA7^+^-DLL1^+^ sister pairs showed marker expression. **h,** Time-lapse images of two neighboring cells of which one expresses *DLL1* (red asterisk) and the other *CA7* (white asterisks). Note that these cells are not sister cells. Dashed white lines indicate the pair of cells at before clear marker expression. Scale bar: 10 µm. **i,** Neighbors of present and future DLL1^+^ cells were evaluated 60 hours after shift to differentiation medium. Null hypothesis that all three cell types (BEST4/CA7^+^, DLL1^+^ and unmarked) had equal probability to have a DLL1^+^ neighbor was tested using Pearson χ^2^-test.

Next, we labeled *DLL1*-expressing cells by knock-in of a *P2A-Clover* cassette at the C-terminus of the *DLL1* gene (*DLL1-P2A-Clover*) in the *CA7-P2A-tdTomato*; *H2B-iRFP670* reporter organoids and tracked cells during differentiation (**Fig. 3c and Extended Data** Fig. 7c,d). The resulting lineages of DLL1^+^ cells showed *DLL1* expression after a median time of 35 hours (**Fig. 3d-f**), thus even earlier than *MUC2* expression after 49 hours, as shown above (**Fig. 1f,g**). This agrees with findings that *DLL1* expression starts low in the crypt and precedes functional marker expression of secretory cells^35^. Notably, out of 25 cells that were tracked from birth and adopted the BEST4/CA7^+^ fate, 13 had a DLL1^+^ sister, each of which expressed *DLL1* before *CA7* emerged in the BEST4/CA7^+^ cell (**Fig. 3g and Extended Data** Fig. 8). Moreover, at 60 hours after the shift to differentiation medium, all present and future BEST4/CA7^+^ cells were in direct contact with DLL1^+^ cells (**Fig. 3h,i**). This suggested that contact with a DLL1^+^ lineage around 60 hours is necessary for committing to BEST4/CA7^+^ fate. Overall, these results show that DLL1^+^ cells, of which the majority is a goblet cell or it’s precursor, instruct commitment to the BEST4/CA7^+^ cell fate by lateral Notch-signaling.

### BEST4/CA7^+^ cells support the survival of goblet cells

Having established that Notch ligand-presenting goblet cells instruct differentiation of BEST4/CA7^+^ fate in neighboring cells, we wondered if there was a functional relevance for this interaction. We and others previously established that the transcription factor *SPIB* is essential for BEST4/CA7^+^ cell differentiation by showing that genetic knock-out of *SPIB* leads to a complete lack of BEST4/CA7^+^ cells^12,22^. Here, we then examined whether BEST4/CA7^+^ cells impacted goblet cell abundance, by comparing goblet cell numbers in both *SPIB*-KO and wild-type (WT) organoids using FACS analysis. Strikingly, goblet cell fractions in *SPIB*-KO and WT organoids were similar up until day 4 of differentiation but rapidly decreased in *SPIB*-KO organoids on day 5 and 6 (**Fig. 4a,b**). To study the survival of goblet cells more directly, we used live-cell imaging and tracking. Tracking showed that goblet cells were significantly more likely to undergo apoptosis in *SPIB*-KO organoids (compared to WT organoids), as evidenced by fragmentation of the nucleus and the mucus granules (**Fig. 4c,d, Extended Data** Fig. 9a,b **and Supplementary Videos 3,4**). These observations thus revealed a novel function for BEST4/CA7^+^ cells, namely, to increase goblet cell survival.

**Fig. 4:**
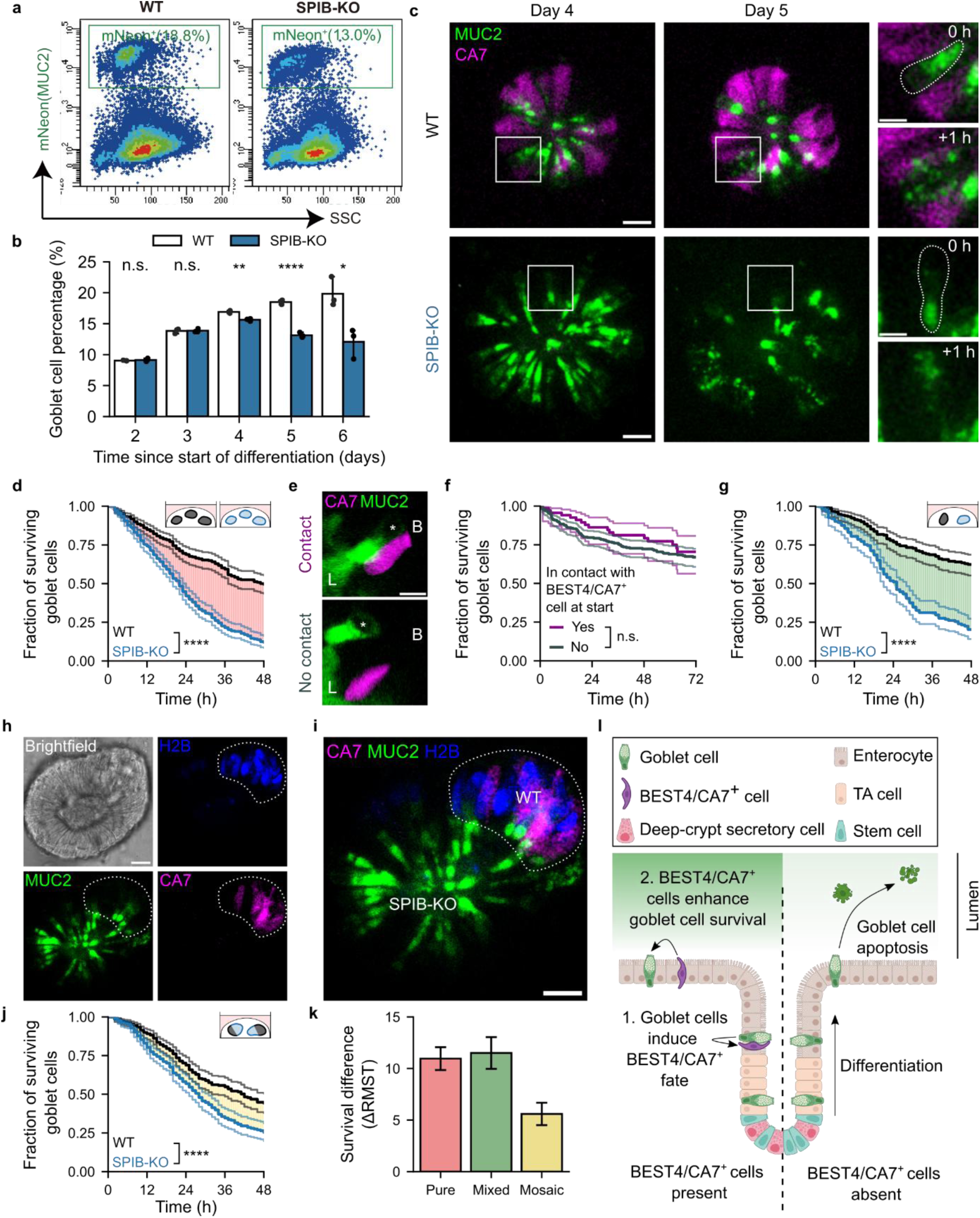
BEST4/CA7^+^ cells enhance goblet cell survival via the lumen. a, Representative FACS plots of goblet cell percentages in WT and *SPIB*-KO organoids after 6 days of differentiation. **b,** Goblet cell percentage on day 2 to day 6 since shift to differentiation medium, as quantified by FACS analysis. n = 3 biological replicates. Student’s t-test is used for comparisons. **** p < 0.0001, ** p < 0.01, * p < 0.05, n.s.: not significant. Error bars indicate 95% confidence interval. **c,** Examples of goblet cell apoptosis in WT (top) and *SPIB*-KO (bottom) organoids. *SPIB*-KO lack BEST4/CA7^+^ cells. Insets show time point of goblet cell death. Scale bars, 20 µm in the large images and 10 µm in the insets. **d,** Goblet cell survival in differentiated organoids. Data are derived from 365 WT goblet cells and 393 *SPIB*-KO goblet cells. N = 16 organoids derived from three independent imaging experiments are analyzed per condition. Logrank test at 48 hours is used for comparison. **** p < 0.0001. **e,** Micrographs showing goblet cells in contact (top) and one not in contact (bottom) with a BEST4/CA7^+^ cell. Asterisk indicates the nucleus of goblet cells. L and B indicate the luminal and basal side of the organoids, respectively. Scale bar: 10 µm. **f,** Survival curves for goblet cells with and without contact with a BEST4/CA7^+^ cell at the start of imaging. Imaging was started after day 6 of differentiation. Data are derived from 69 goblet cells with and 290 goblet cells without contact with a BEST4/CA7^+^ cell. N = 28 organoids derived from three independent imaging experiments are analyzed. Logrank test at 72 hours is used for comparison, n.s.: not significant. **g,** Survival curves for goblet cells in co-cultures of WT and *SPIB*-KO organoids. Data are derived from 210 WT goblet cells and 162 *SPIB*-KO goblet cells in mixed cultures. N = 15 organoids derived from two independent imaging experiments are analyzed. Logrank test at 48 hours is used for comparison. **** p < 0.0001. **h,i,** Micrographs of a WT/*SPIB*-KO mosaic organoid. White dotted lines indicate the part of the organoid containing WT cells (with nuclear H2B-iRFP670 florescence). The rest of the organoid consists of *SPIB*-KO (H2B-iRFP670^-^) cells. Scale bars: 20 µm. **j,** Survival curves of *SPIB*-KO and WT goblet cells in mosaic organoids. Data are derived from 363 WT goblet cells and 412 *SPIB*-KO goblet cells. N = 23 organoids derived from three independent experiments are analyzed. Logrank test at 48 hours is used for comparison. **** p < 0.0001. Confidence interval for survival curves in panels **(d)**, **(f)**, **(g)** and **(j)** are computed using Greenwood’s exponential formula. **k,** Difference in survival between WT and *SPIB*-KO goblet cells in pure cultures, mixed cultures and mosaic cultures, measured as the difference in Restricted Mean Survival Times (ΔRMST) up to t = 48 hours. ΔRMST corresponds to the area between the survival curves, as shown with red, green and yellow shaded areas in panels **(d)**, **(g)** and **(j)**. Error bars indicate the standard error (SE) of the ΔRMST. **l,** Proposed model for interdependence of goblet and BEST4/CA7^+^ cells in the human intestinal epithelium.

To understand how goblet cell survival is supported by BEST4/CA7^+^ cells, we tested whether it depended on direct cell-cell contact or was instead mediated indirectly – either via the lumen (apically) or the medium (basally). In WT organoids, survival of goblet cells beyond day 6 did not depend on BEST4/CA7^+^ cell contact (**Fig. 4e,f and Extended Data** Fig. 9c). The difference in goblet survival observed above between WT and *SPIB*-KO organoids was not decreased by co-culturing these organoids in a 1:1 ratio (**Fig. 4g and Extended Data** Fig. 9d), arguing against signals secreted basally into the shared medium. To determine whether goblet cell survival required a shared lumen between the goblet and BEST4/CA7^+^ cells, we assessed goblet cell survival in mosaic organoids consisting of *SPIB*-KO and WT cells (with WT cells labeled by H2B-iRFP670, **Fig. 4h,i**). Notably, the survival of SPIB-KO goblet cells was much closer to that of the WT goblet cells in the same mosaic organoid (**Fig. 4j,k**), as compared to the previous two conditions (**Fig. 4g-i**). Together, these results show that goblet cell survival beyond day 4 of differentiation does not rely on direct cell-cell contact but instead depends on the shared lumen.

## Discussion

By quantifying spatial-temporal differentiation dynamics using live-cell imaging, we uncovered a cell abundance-control mechanism based on an interdependence between BEST4/CA7^+^ and goblet cells (**Fig. 4l**). Goblet cells – as the primary source of Notch-activating DLL1 –laterally activate Notch^30,31^ signaling in their sister cells, as well as in other direct neighbors, thereby instructing BEST4/CA7^+^ cell fate commitment in these neighbors. In turn, the long-term survival of goblet cells requires the presence of BEST4/CA7^+^ cells. The latter interaction does not depend on direct contact and is mediated by their shared lumen. Thus, goblet cells induce BEST4/CA7^+^ cells that in turn support their long-term survival.

The existence of reciprocal relations between intestinal epithelial cells that control not only differentiation but also survival is notable. Interactions between different cell types in the intestinal epithelium are well-known to drive differentiation and thereby contribute to proper homeostasis. Paneth cells are known to provide EGF, Wnt and Notch ligands that help to maintain stemness^36^. Tuft cells were shown to increase goblet and tuft cell numbers during helminth infections^37,38^, though this interaction was not strictly intra-epithelial, depending on secretion of cytokines by innate lymphoid cells. M cells were shown to inhibit further M cell differentiation by lateral inhibition^39^ and expression of RANKL decoy receptors^40^. Yet, cell abundance is in principle equally dependent on cell survival, which we show here is regulated by intra-epithelial cellular interactions. Such reciprocal cooperation between cells may occur more generally and be important to maintaining and adapting the intestinal barrier during normal and inflammatory conditions.

The observed reciprocal interactions may impact medical intervention as well. Pharmacological blocking of Notch-ligands DLL1 and DLL4 has been explored as a strategy for reducing tumor growth^41^. This is based partly on insights from the mouse intestinal epithelium, where BEST4/CA7^+^ cells are absent^16^, and blocking Dll1 moderately increases goblet abundance without the complete loss of any cell type^42,43^. However, here we show that in the human intestinal epithelium, abrogated DLL1 expression leads to an almost complete loss of BEST4/CA7^+^ cells, which in turns causes strongly enhanced goblet cell death. Targeting DLL1 in the human intestine will therefore likely generate responses that differ strongly from those in the mouse. Our findings thus highlight the important differences between human and mouse intestinal epithelia.

The interdependence between BEST4/CA7^+^ and goblet cells may be particularly relevant for conditions such as Crohn’s disease and ulcerative colitis, which are associated with goblet cell depletion^17–21^. Interestingly, the expression of mature BEST4/CA7^+^ marker genes in the colon are then also substantially decreased^4,9,44^ (**Extended Data** Fig. 10). Decreasing abundance of BEST4/CA7^+^ cells may thus limit goblet cell survival in these diseases, while goblet cell depletion may conversely also cause BEST4/CA7^+^ cell number decline. This model is consistent with earlier reports showing that goblet cells are specifically depleted in the upper part of the crypt regions of patients with active ulcerative colitis^17^. The results provide new perspectives on identifying the upstream causes of goblet and BEST4/CA7^+^ cell depletion, dependence on signals detected by goblet and BEST4/CA7^+^ cells, downstream effects on epithelial integrity, and the design of alternative therapeutic interventions.

## Online Methods

### Human intestinal organoid lines

Human colon organoids were established previously in our lab^45^. The colon tissue (Donor No: P11N) was obtained from the Diakonessen Hospital Utrecht, with informed consent. The study was approved by the ethical committee and was conducted in accordance with the Declaration of Helsinki and Dutch law. This study complied with all relevant ethical regulations regarding research involving human participants.

### Organoid culture

Organoids were cultured as previously described^12,46^. Briefly, organoids were embedded in small 5-10 µL droplets of Cultrex Basement Membrane Extract, growth factor reduced type 2 (BME2; R&D Systems; cat. #3536-005-02) and kept in expansion medium. Expansion medium consists of adDMEM/F12 (Gibco; cat. #12634028) supplemented with 100 U/ml Penicillin/Streptomycin (P/S; Gibco; cat. #15140122), 10 mM HEPES (Gibco; cat. #15630056), 1× Glutamax (Gibco; cat. #35050038), 1× B-27 supplement (Thermo Fisher; cat. #12587010), 1.25 mM N-acetyl-L-cysteine (NAC; Sigma-Aldrich; cat. #A9165), 1% (v/v) recombinant Noggin in conditioned medium (U-Protein Express BV; custom order), 0.5 nM WNT surrogate (U-Protein Express BV; Custom order), 50 ng/ml human Epidermal Growth Factor (EGF; Peprotech; cat. #AF-100-15), 0.5 μM A83-01 (Tocris; cat. #2939), 1 μM SB202190 (Sigma-Aldrich; cat. #S7067), 1 μM Prostaglandin E2 (PGE2; Tocris; cat. #2296), 10 mM Nicotinaminde (NIC; Sigma-Aldrich; cat. #N0636) and 20% (v/v) recombinant R-spondin 1 in conditioned medium (U-Protein Express BV; custom order).

To differentiate organoids, expansion medium was removed, and cells were incubated for 2 hours in adDMEM/F12 supplemented with 100 U/ml P/S, 10 mM HEPES and 1× Glutamax (wash medium) for 2 hours. Then, differentiation medium was added, which had the same composition as expansion medium except that EGF, Noggin, SB202190, A83-01, WNT surrogate, PGE2 and NIC.

### Generation of genetically modified organoids

Generation of *CA7-P2A-tdTomato* and *MUC2-mNeonGreen* reporter organoids, by CRISPR-HOT approach, has been described in our previous study^12^. *DLL1-P2A-Clover* knock-in reporter organoids were similarly generated using CRISPR-HOT^24,47^. Three plasmids were used: (1) the frame-selector plasmid containing an sgRNA (to linearize the donor-targeting vector), Cas9 and mCherry (for the detection and FACS sorting of the successfully transfected cells) was obtained from Addgene (plasmid# 66941); (2) The *P2A-Clover* donor-targeting vector was obtained from Addgene (plasmid# 138568) and (3) the sgRNA vector (Addgene, plasmid# 47108) containing sequence targeting 3’ end of *DLL1* gene. The sgRNA sequence is 5’-TGAGTGCGTCATAGCAACTG-3’. 5 μg sgRNA vector, 5 μg frame-selector plasmid and 5 μg donor-targeting vector were co-transfected into organoid cells using the NEPA electroporation system (NEPAGENE). After FACS sorting, based on the mCherry fluorescence, subclones were picked and expanded in expansion medium. Successful insertions were first identified by direct visualization of the fluorescence marker in the differentiated organoids and then confirmed by targeted genotyping via Sanger sequencing.

The H2B-iRFP670 sequence was integrated into the genome using mT2TP transposase system. 5 μg transposase plasmid and 5 μg donor plasmid with terminal inverted repeats (TIRs) bearing H2B-iRFP670 sequence were co-transfected into organoid cells by electroporation.

Generation of *SPIB* knock-out organoids has been described in our previous study^12^. *DLL1* and *DLL4* knock-out organoids were similarly generated by introducing an early stop codon) using CRISPR C-to-T base-editing: 7.5 μg spCas9-CBE6b plasmid (Addgene, plasmid# 215820), 2.5 μg sgRNA plasmid and a two-plasmid transposon system (5 μg PiggyBac transposase plasmid + 5 μg donor plasmid with terminal inverted repeats (TIRs) bearing hygromycin resistance for organoid selection) were co-transfected into organoid cells through electroporation. After FACS sorting of DAPI^-^ live single cells followed by hygromycin selection, subclones were picked and expanded in expansion medium. Successful homozygous knock-out organoids were confirmed by targeted genotyping via Sanger sequencing. The sgRNA sequence for *DLL1*-KO and *DLL4*- KO are: 5’-TCCCAGGACCTGCACAGCAG-3’ and 5’-CATCCAGGGCTCCCTAGCTG-3’, respectively.

Genotyping primers used for knock-in and knock-out validation were listed in **Supplementary Table 1**.

### Sample preparation and flow cytometry

Organoids were released from BME using ice-cold Corning Cell Recovery Solution (Sigma-Aldrich; cat. #CLS354253) and dissociated with 1 ml TrypLE^TM^ Express Enzyme (TrypLE; Thermo Fisher; cat. #12605010) at 37 °C for 6-8 mins, followed by gently pipetting 20 times. After TrypLE dissociation, the cell suspension was filtered through a 40 µm cell strainer and stained with DAPI (Sigma-Aldrich; cat. #10236276001) for FACS analysis or cell sorting. Samples were analyzed on a BD LSR Fortessa X20 equipped with 4 lasers (BD Bioscience). At least 5,000 DAPI^-^ live cells were recorded for each analyzed sample.

### RNA extraction and qPCR

Organoids were subjected to RNA isolation using a NucleoSpin RNA kit (Macherey-Nagel; cat. #740955.50) following the manufacturer’s protocol. Reverse transcription reactions were performed using GoScript^TM^ reverse transcriptase kit (Promega; cat. #A5000). cDNA was subjected to qPCR analysis using iQ^TM^ SYBR Green Supermix (BioRad; cat. #1708887) on a CFX Connect Real-Time PCR machine (BioRad). For gene expression analysis, qPCR was performed with gene-specific qPCR primers. Ct value of each gene was normalized to GAPDH (as the ΔCt), and fold change between experimental groups was calculated with the 2^-ΔΔCt^ method. All qPCR primers used in this study are listed in **Supplementary Table 1**.

### Long-term live-cell imaging

For tracking experiments of differentiating and expanding organoids, organoids were dissociated^46^ and plated onto an imaging chamber (CellVis) and expanded for 2-3 days. Organoids were then washed with wash medium for 2 hours before differentiation medium added. Cells were imaged using a A1R MP (Nikon) scanning confocal microscope with a 1.30 NA 40× magnification oil immersion objective. Images were taken every 12 minutes, and at each time point, 31 Z-slices were imaged with 2 µm intervals.

For tracking goblet cell survival, differentiated live reporter organoids were staged on the Leica SP8 confocal detection system fitted on a Leica DMi8 microscope or a Leica TCS SP8 to capture time series images every 15 – 60 mins for 18 – 48 hours. All tracking experiments were performed at 37 °C and 5% CO2.

### Cell tracking

Tracking of expansion and differentiation of human colon organoids was done manually using our custom tracking software OrganoidTracker^25^. For tracking the organoids in expansion medium, we selected three organoids and tracked the nuclei from the bottom 10 µm of the organoid at the start, and continued tracking until cells extruded, moved out of view, or were no longer trackable with high fidelity.

For tracking the remaining differentiating organoids, we tracked We reconstructed the full lineage tree of all observable CA7^+^, MUC2^+^ and DLL1^+^ cells. Organoids were excluded from this analysis if they moved out of view, collapsed, contained CA7^+^, MUC2^+^ or DLL1^+^ cells already at the start of imaging, or contained no cells expressing *CA7*, *MUC2* or *DLL1* throughout the entire time-lapse.

### Single-cell RNA sequencing analysis

Single-cell RNA sequencing data of the human colon epithelium was retrieved from the GutCellAtlas^6,7^. The .h5ad file was loaded as an AnnData object and processed using the scanpy library. Only cells with less than 20% mitochondrial gene expression, derived from primary healthy pediatric and adult human colon epithelium were included. Furthermore, only genes expressed in at least three cells were included in the analysis. Cell types with less than 20 cells were excluded from this analysis. Data was normalized (target sum 10.000), log10 transformed, and the top 2,000 highly variable genes were found. Data was then scaled, and 20 principal components (PCs) coordinates were found, and the nearest neighbor distance matrix and neighborhood graph was computed (knn = 15) using all 20 principal components. Then, dimension reduction was performed using t-distributed stochastic neighborhood embedding (tSNE) using 20 components.

### Mosaic organoids

To generate the mosaic organoids, WT (labeled with H2B-iRFP670) and *SPIB*-KO double reporter organoids were dissociated into single cells (as described above), mixed at a 1:1 ratio, and plated into a BME-coated transwell insert or culturing well for 2D attachment for 2 days in the expansion medium. Then, the monolayer of mixed cells was dissociated, seeded as BME droplets, and cultured in expansion medium for organoid growth.

### Data retrieval from IBD TaMMA database

Expression levels of selected proteins in colon tissue samples of patients with Crohn’s Disease (CD), Ulcerative Colitis (UC) and healthy control were retrieved from the IBD Transcriptome and Metatranscriptome Meta-Analysis (IBD TaMMA) database^44^. P-values of statistical tests for proteins of interest were obtained from the IBD TaMMA web-app (https://ibd-meta-analysis.herokuapp.com/).

### Statistics

Statistical analyses were performed as indicated in the figure legends. Mann-Whitney U test, Student’s t-tests, Pearson χ^2^-tests were performed using the scipy.stats library in Python^48^, with the following significance: **** p < 0.0001, *** p < 0.001, ** p < 0.01, * p < 0.05, and n.s. p > 0.05. Logrank test and survival analysis were performed using the lifelines library in Python^49^.

Restricted Mean Survival Time (RMST) and corresponding variance were computed directly from the Kaplan-Meier estimates for the survival functions. For given survival curve 𝑆(𝑡), and time horizon 𝑡^∗^, RMST is computed via

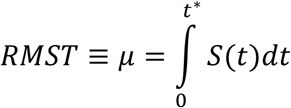

Given two survival curves 𝑆_0_(𝑡) and 𝑆_1_(𝑡), the RMST difference (Δ) corresponds to the area between survival curves and serves as a measure of the effect size for the difference in survival between two groups even if the proportional hazard assumption is breached^50^. The Restricted Standard Deviation of Survival Time (RSDST), which corresponds to the standard deviation of the RMST, was computed as^50^:

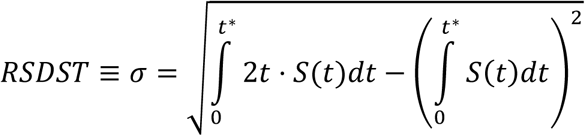

For each estimated RMST difference (Δ^ = 𝜇^_1_ − 𝜇^_0_) between two groups with Kaplan-Meier survival curves 𝑆^_0_(𝑡), 𝑆^_1_(𝑡), sample sizes 𝑛_0_, 𝑛_1_ and RSDST estimates 𝜎^_0_, 𝜎^_1_ , the standard error was computed as

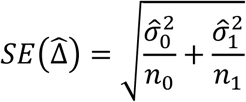

### Software

Image processing was done in Fiji/ImageJ (v1.54g). Tracking was performed in OrganoidTracker^25^. FACS data was plotted using BD FACS Diva software (v8.0.1), FlowJo (v10). Downstream data analysis and plotting was done in Python 3.9, with the use of the following libraries: numpy (v1.26.0, scipy (v1.12.0), pandas (v2.1.0), seaborn (v0.13.0), matplotlib (v3.6.3), lifelines (v0.27.8), scanpy (v1.9.3). Figures were made in Inkscape (v1.1.1; 3bf5ae0d25, 2021-09-20).

## Author contributions

Conceptualization: all authors collaboratively; Methodology: W.K.S. and D.W.; Software: W.K.S.; Validation: W.K.S. and D.W.; Formal analysis: W.K.S. and D.W.; Investigation: W.K.S. and D.W.; Resources: D.W.; Data Curation: W.K.S.; Writing – original draft: W.K.S.; Writing – Review and Editing: W.K.S., D.W., S.J.T., J.v.Z., H.C.; Visualization: W.K.S. and D.W.; Supervision: S.J.T., J.v.Z., J.H.v.E., H.C.; Project and administration: all authors; Funding acquisition: J.v.Z., S.J.T., J.H.v.E., H.C.

## Acknowledgements

We thank all members of the Clevers, van Zon and Tans labs for fruitful discussions. We thank Marko Kamp, Corinna Hoefler, Yvonne Goos for technical support. This work is supported by the project Organoids in Time with project no. 2019.085 of the research program NWO Investment Large financed by the Dutch Research Council (to J.S.v.Z., S.J.T. and H.C.), the Netherlands Organ-on-Chip Initiative, an NWO Gravitation project (no. 024.003.001) funded by the Ministry of Education, Culture and Science of the Government of the Netherlands (to H.C.).

## Competing interests

H.C. is head of Pharma Research and Early Development (pRED) at Roche. H.C. is inventor of several patents related to organoid technology; his full disclosure is given at https://www.uu.nl/staff/JCClevers/.

## Data availability

Further information and requests for resources should be directed to Sander J. Tans. Requests for human organoid lines should be directed to Hans Clevers. Sharing the organoid lines used in this study requires approval from local Institutional Review Board. A complete Materials Transfer Agreement will also be required. This study didn’t generate original sequencing datasets.

## Code availability

OrganoidTracker source code is publicly available at the OrganoidTracker Github page (https://github.com/jvzonlab/OrganoidTracker).

## Supplementary Note 1: Inclusion criterion for unmarked cells

For quantification of the frequencies of sister pairs, every cell type must be assigned either BEST4/CA7^+^ cell, goblet cell or unmarked cell fate. Cells are, however, frequently lost from the imaging dataset, either due to cell death or because the cell moves out of the imaging window. It is therefore possible that some cells without BEST4/CA7^+^ or goblet cell marker when they are lost from the dataset would have shown the markers later. To avoid overrepresentation of Unmarked cells in the dataset, we filtered out unmarked cells for which the probability was > 5% that they would have expressed the BEST4/CA7^+^ or goblet cell marker if they could have been tracked for longer (**Extended Data** Fig. 4a). This amounts to excluding cells that were lost from the imaging dataset before the point in time, denoted as 𝑡_0.95_, when a given cell still has a > 5% probability to express either marker.

Let 𝑈_𝑡_, 𝐺_𝑡_and 𝐵_𝑡_be the events that a cell is an unmarked cell, a goblet cell or a BEST4/CA7^+^ cell at time 𝑡, respectively. The probability that a cell would have remained unmarked (event 𝑈_∞_), given that it is unmarked at time 𝑡, is:

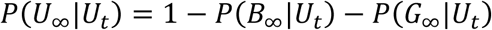

Using Bayes’ theorem, this can be rewritten as:

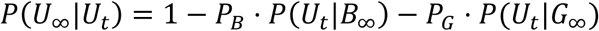

Here 𝑃_𝐵_and 𝑃_𝐺_are the a priori probabilities (base-rates) of cells committing to goblet or BEST4/CA7^+^ cell fate, which we previously found to be 𝑃_𝐵_ ≈ 1% and 𝑃_𝐺_ ≈ 19% using FACS- sorting^12^. The remaining conditional probabilities, which represent the probabilities that an eventual BEST4/CA7^+^ or goblet cell is still unmarked at time 𝑡 can be estimated from the probability density functions of the marker expression times (**Fig. 1f and Extended Data** Fig. 4b). We approximate these distributions as normal distributions, so that:

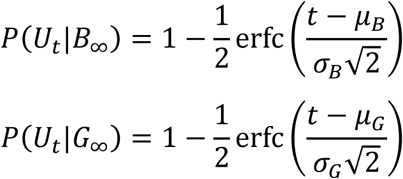

Here erfc(𝑥) = 1 − erf (𝑥) is the complement of the error function, the constants 𝜇_𝐵_, 𝜎_𝐵_ and 𝜇_𝐺_, 𝜎_𝐺_ indicate the mean and standard deviations of marker expression times for BEST4/CA7^+^ and goblet cells, respectively. Together, this means that the sought probability is given by:

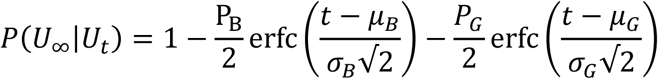

By equating this probability to 95% and solving for 𝑡 , we find that 𝑡_0.95_ = 69.6 hours (**Extended Data** Fig. 4c).

## Supplementary Note 2: Computing of baseline expectation of sister pairs

Here, we derive the equations used for computing the expected numbers of distinct sister pairs under the baseline assumption that cell fate commitment to either the BEST4/CA7^+^ (B), goblet (G) or unmarked (U) fate is fully cell-autonomous and independent of lineage history and spatiotemporal cues. In this picture, the three cell types B, G and U are randomly distributed over lineages, with the probability that two cells of particular type are sisters given by the product of the frequencies at which each type occurs within the population.

Let 𝑃_𝐵_ and 𝑃_𝐺_be the population level probabilities (priors) that a given cell commits to the BEST4/CA7^+^ and goblet cells. Using FACS-experiments, 𝑃_𝐵_ ≈ 1% and 𝑃_𝐺_ ≈ 19% in previous work^12^ (see also **Note S1**). In general, the set of possible sister pairs (𝑆) consists of 9 possible sister pairs, namely:

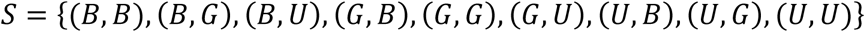

In our dataset, however, we only tracked lineage trees of lineages containing at least one BEST4/CA7^+^ or goblet cell and therefore excludes 𝑈𝑈 sister pairs. Thus, our dataset 𝐷 ⊂ 𝑆 contains the following elements:

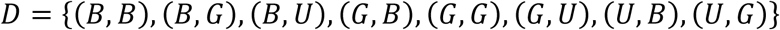

The probability that a given sister pair in our dataset 𝐷 is of type (𝑖, 𝑗) under the baseline assumption is then given by:

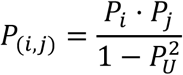

Here the denominator 1 − 𝑃^2^ ensures that the probabilities that a sister pair is of type (𝑖, 𝑗) within our dataset 𝐷 sum up to 1.

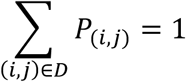

Noting that 𝑃_𝑈_ = 1 − 𝑃_𝐵_ − 𝑃_𝐺_ and therefore 𝑃_𝑈𝑈_ = (1 − 𝑃_𝐵_ − 𝑃_𝐺_)^2^, we obtain 𝑃_𝑖_ ⋅ 𝑃_𝑗_

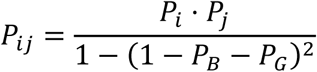

Since we do not distinguish between the order of the cell types in asymmetric sister pairs, we are interested in the probabilities of each possible sister pair independent of the order. That is, we look at the symmetrized dataset 𝐷̅, being:

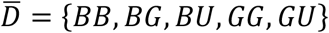

We note that we use the convention to abbreviate the types of sister pairs in alphabetical order, as BB, BG, BU, GG and GU.

Now, the probability of each possible sister pair 𝑖𝑗 ∈ 𝐷̅ is given by:

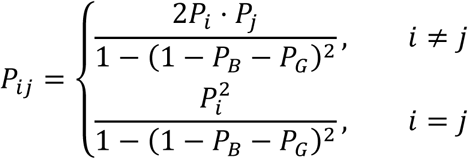

The expected numbers of sister pairs is then calculated using:

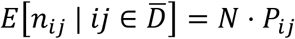

## Extended Data

**Extended Data Fig. 1:**
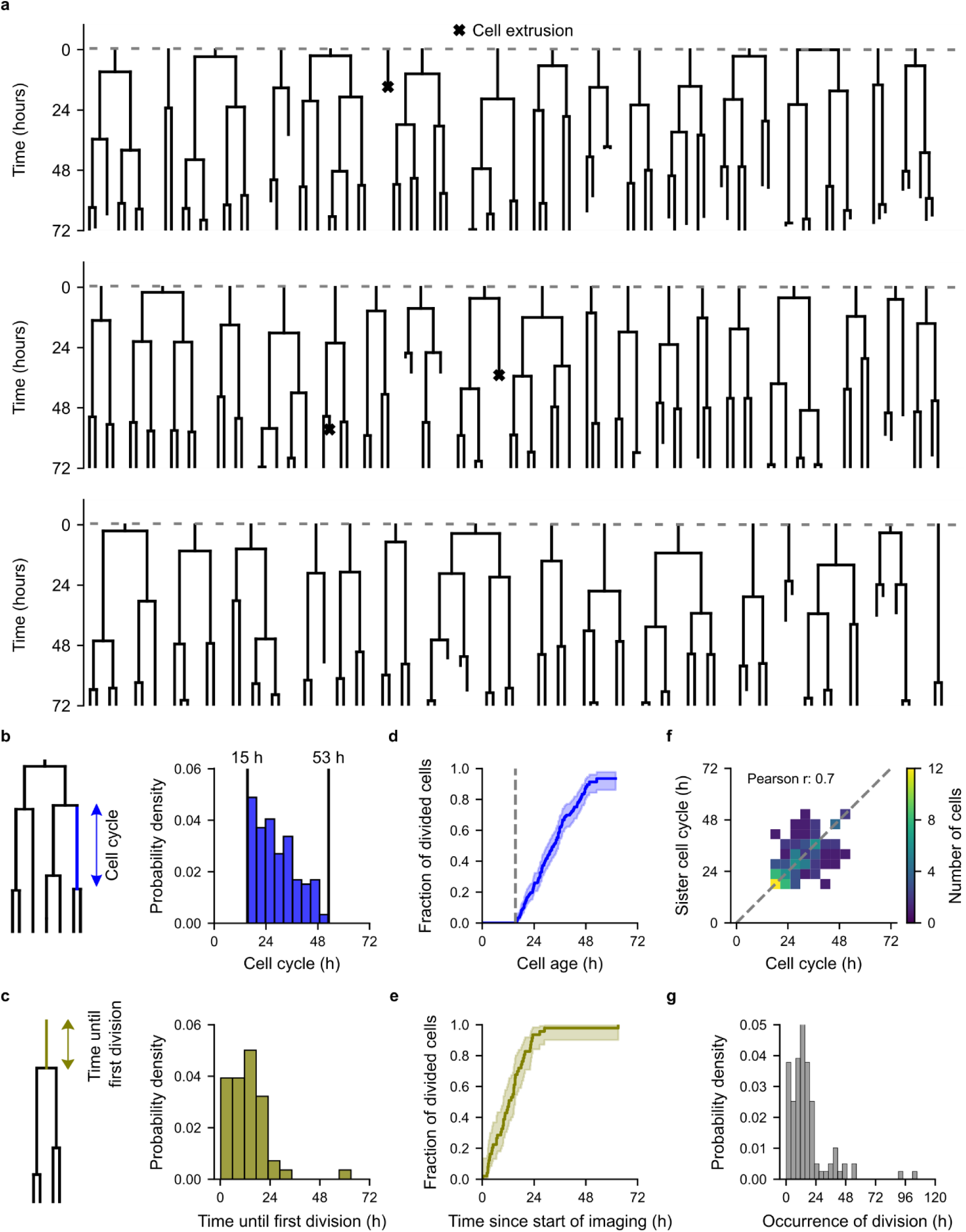
Expansion medium keeps cells in a proliferative and undifferentiated state. a, Lineage tree gallery of three organoids in expansion medium. b,c, Distribution of cell cycle durations (b) and time until division of cells present at the start of imaging (c). d,e, Cumulative distribution of divided cells since their birth (d) and since the start of imaging (e). Error around the distributions due to censoring (e.g. cells moving out of view or extruding) are estimated using a Kaplan-Meier model. Vertical dashed line in (e) indicates the shortest observed cell cycle. f, Correlation of cell cycle durations between sister cells (n = 118 sister pairs). g, Divisions virtually stop occurring after approximately 48 hours in differentiation medium.

**Extended Data Fig. 2:**
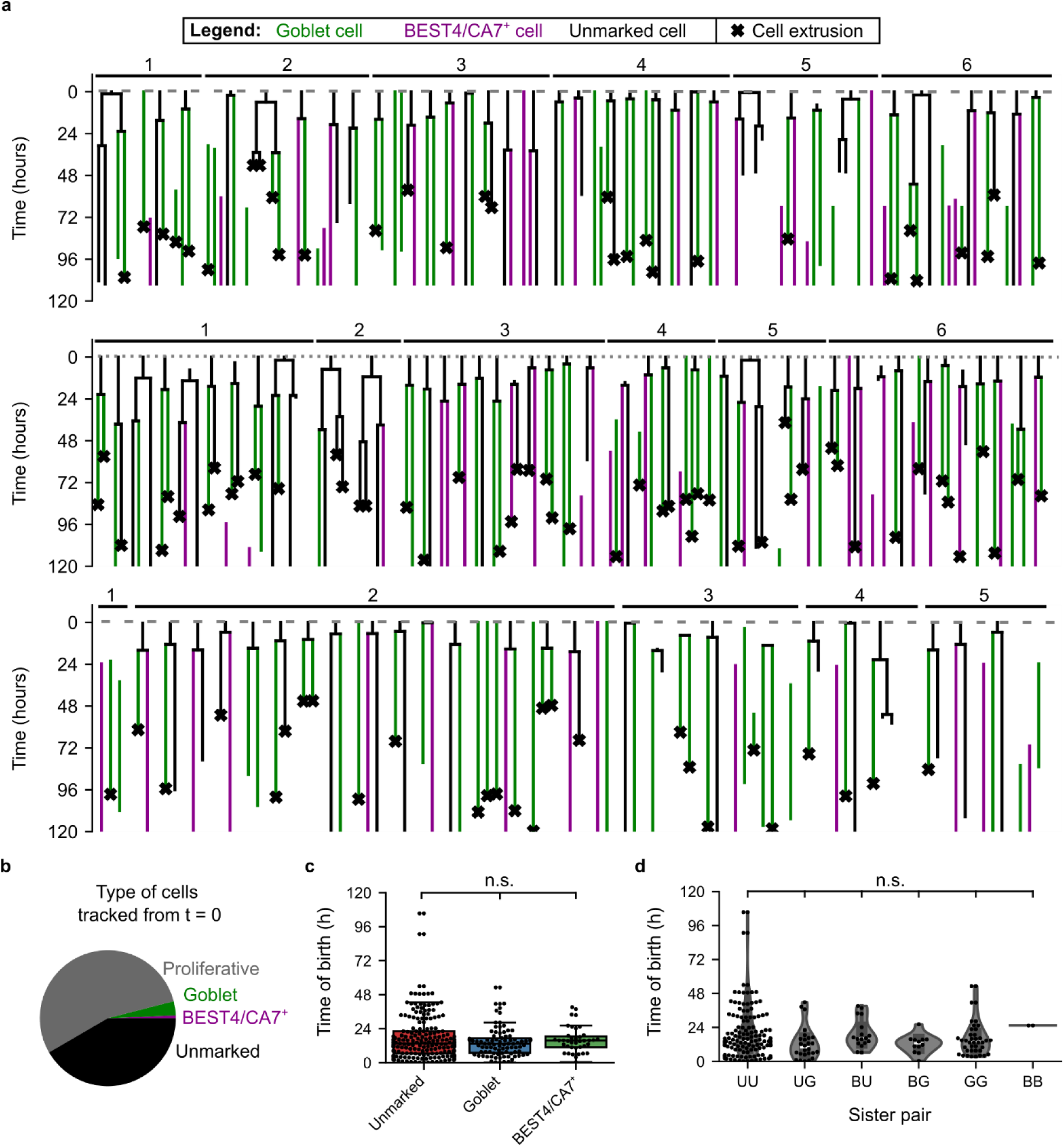
Lineage tree reconstruction of goblet and BEST4/CA7^+^ cells in differentiating organoids. a, Lineage tree gallery of n = 17 organoids derived from three independent tracking experiments. Shown are only lineage trees that contained at least one BEST4/CA7^+^ or goblet cell. b, Identity of the cells present already at time zero. c, Time of birth of unmarked, goblet and BEST4/CA7^+^ cells. Kruskal-Wallis test is used for multiple comparisons, n.s.: not significant. In box plots, the center line shows the median, the box spans the interquartile range (IQR), and the whiskers extend to the most extreme data points within 1.5×IQR from the quartiles. d, Distribution of birth-times for all sister pairs. Kruskal-Wallis test is used for multiple comparisons, n.s.: not significant. Panels (c) and (d) only included cells that were tracked beyond 63.7 hours after the start of differentiation, see Supplementary Note 1.

**Extended Data Fig. 3:**
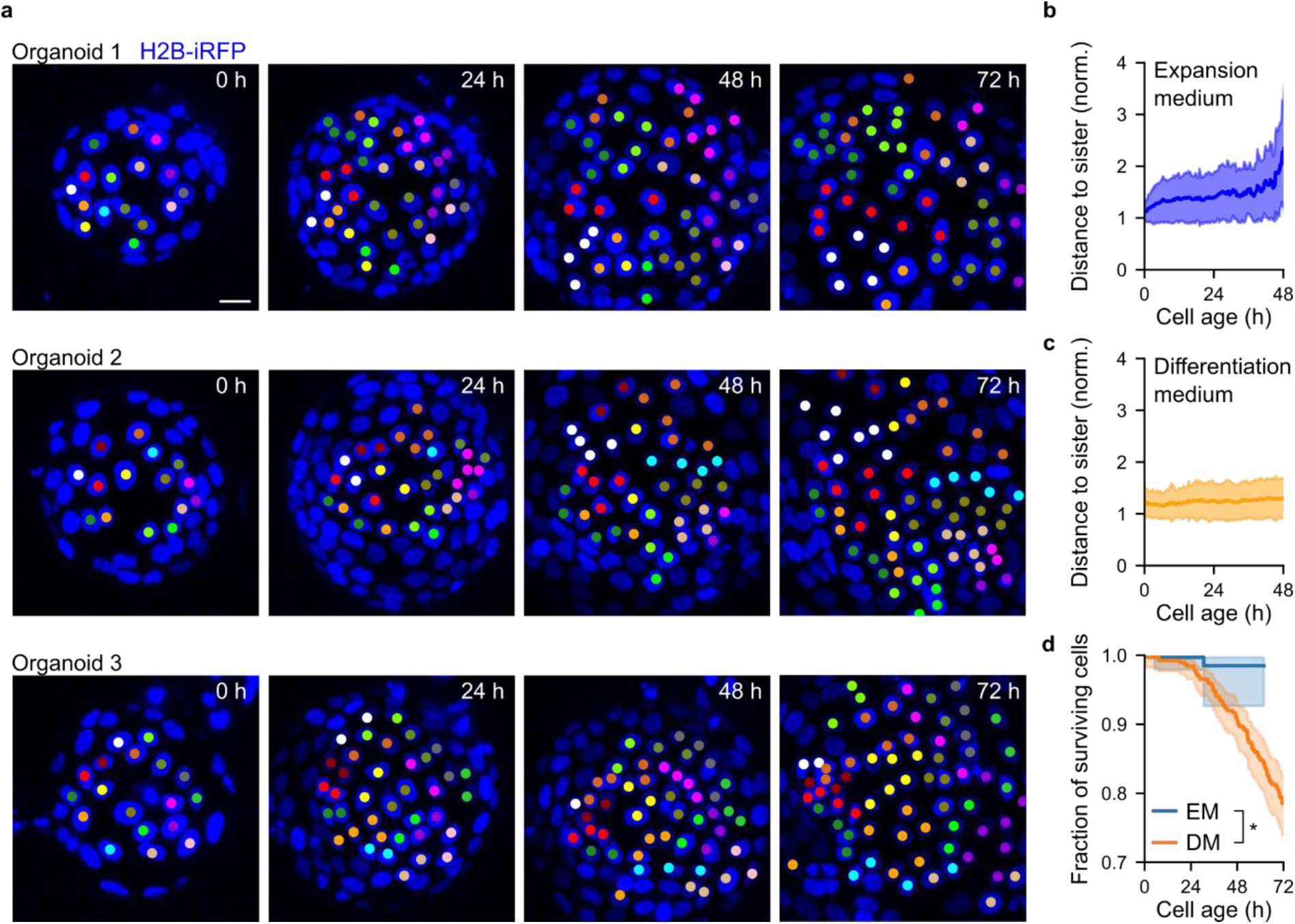
Cell-cell rearrangement rate in the organoids is low in both expansion and differentiation. a, Spatial distribution of clones for three organoids in expansion medium. The color of the dot in the nuclei indicates from which cell the given cell has descended. b,c, Distance to sister cell normalized by the distance to the closest cell for organoids in expansion medium (b) and differentiation medium (c). n = 190 sister pairs (b) and n = 104 sister pairs (c) derived from three organoids in each medium. Only fully tracked organoids were included in (c). Shared area in panels indicates the standard deviation. d, Survival curves of cells in both expansion and differentiation medium estimated from Kaplan-Meier fits. Logrank test is used for comparison at t = 48 h. * p < 0.05. Confidence interval for survival curves are computed using Greenwood’s exponential formula. EM, expansion medium; DM, differentiation medium.

**Extended Data Fig. 4:**
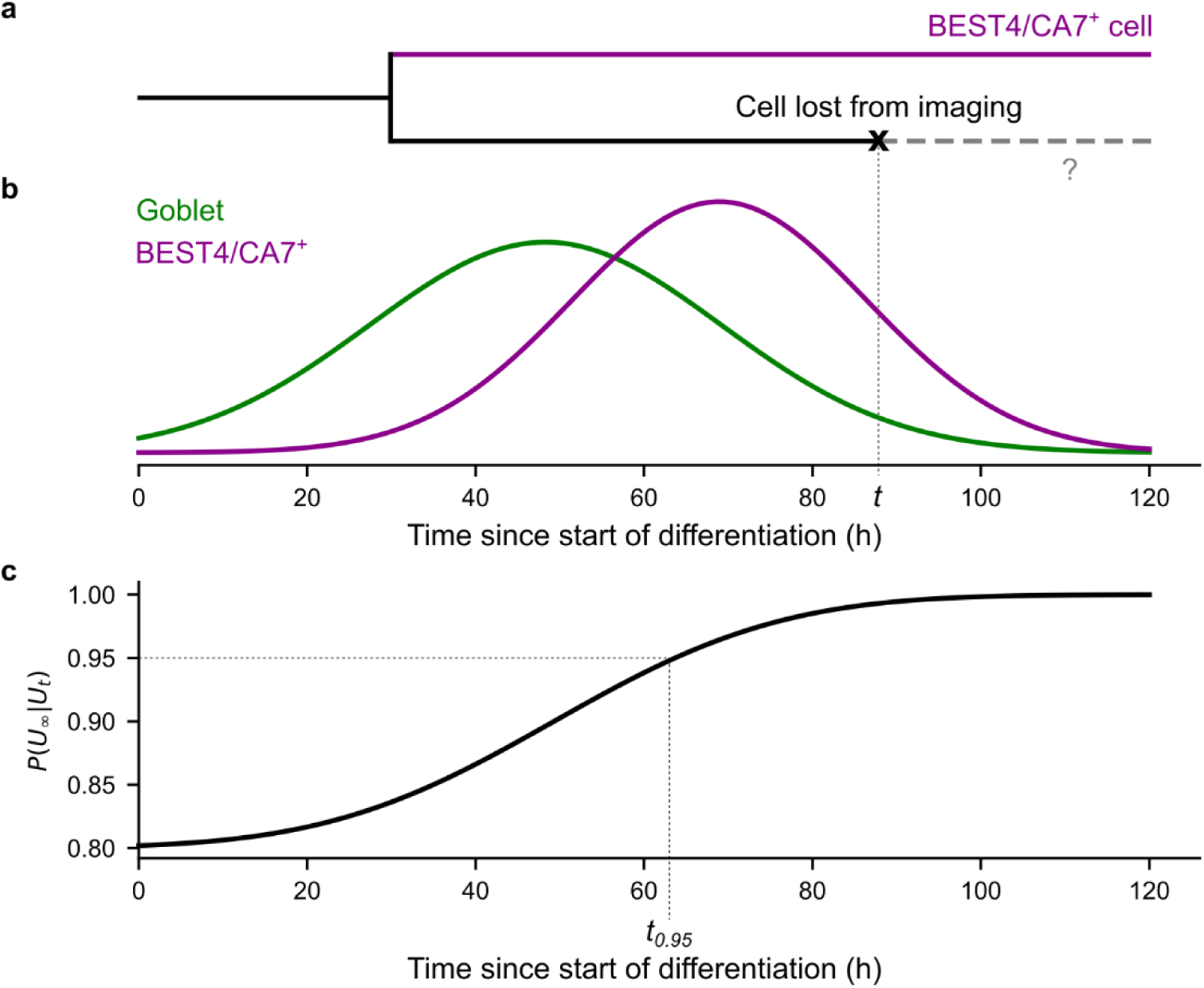
Estimation of the probability that unmarked cells at a given time remain unmarked. a, Hypothetical example of a case where a BEST4/CA7^+^ was tracked until the end of the time-lapse, but its sister was lost from the tracking dataset (either due to death or after moving out of the imaging field) at time 𝑡. b, Probability density functions for marker expression times. c, Probability that an unmarked cell would have remained unmarked as a function of time.

**Extended Data Fig. 5:**
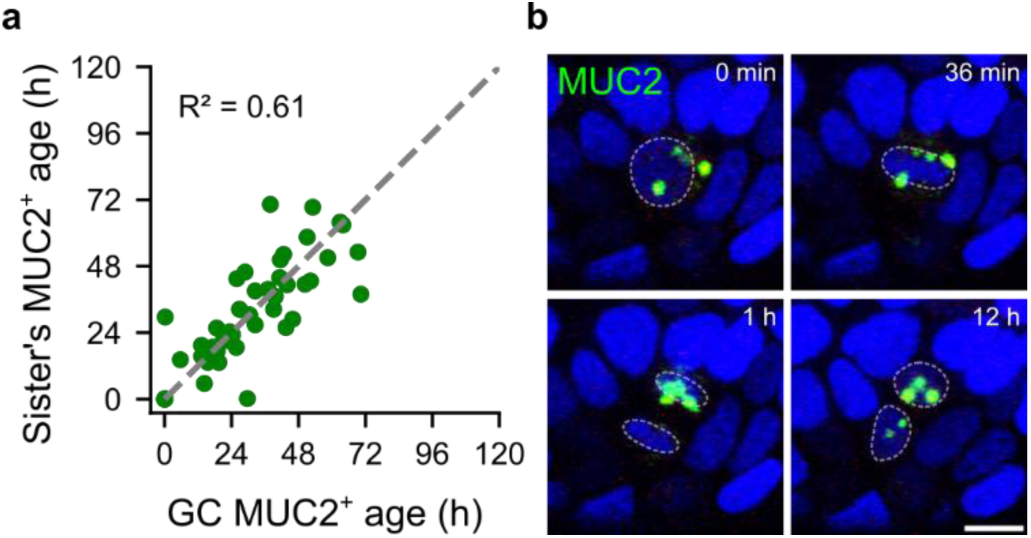
Goblet-goblet sister pairs can be explained by commitment in the mother cell. a, Correlation between age of *MUC2* expression for goblet-goblet cell sisters. R^2^ indicates the coefficient of determination. b, MUC2^+^ proliferative cell dividing into two goblet cells. Scale bar, 10 µm.

**Extended Data Fig. 6:**
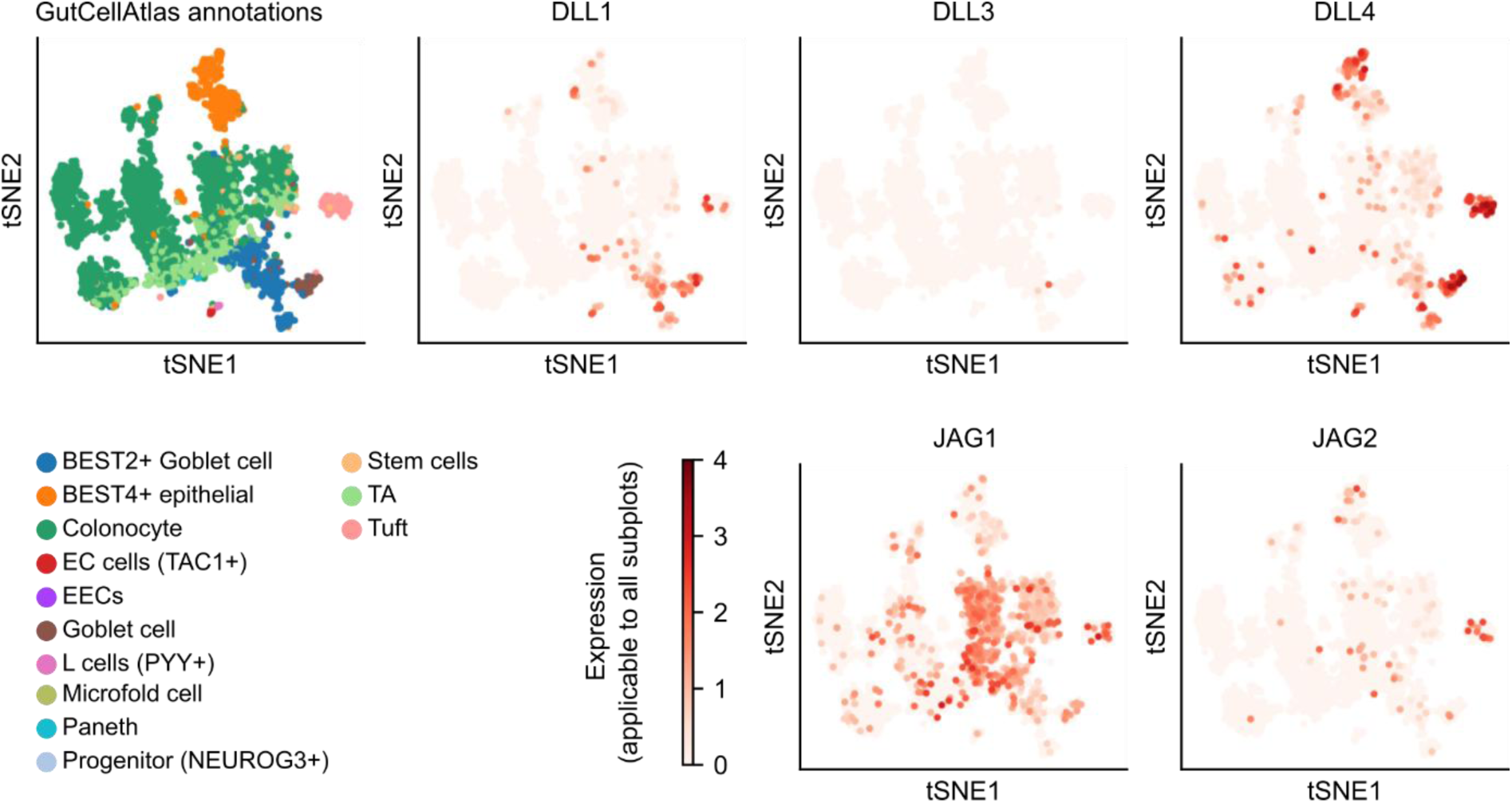
Expression of Notch-ligands in primary human colon epithelial cells. Data and cell type annotations was retrieved from the GutCellAtlas^6^.

**Extended Data Fig. 7:**
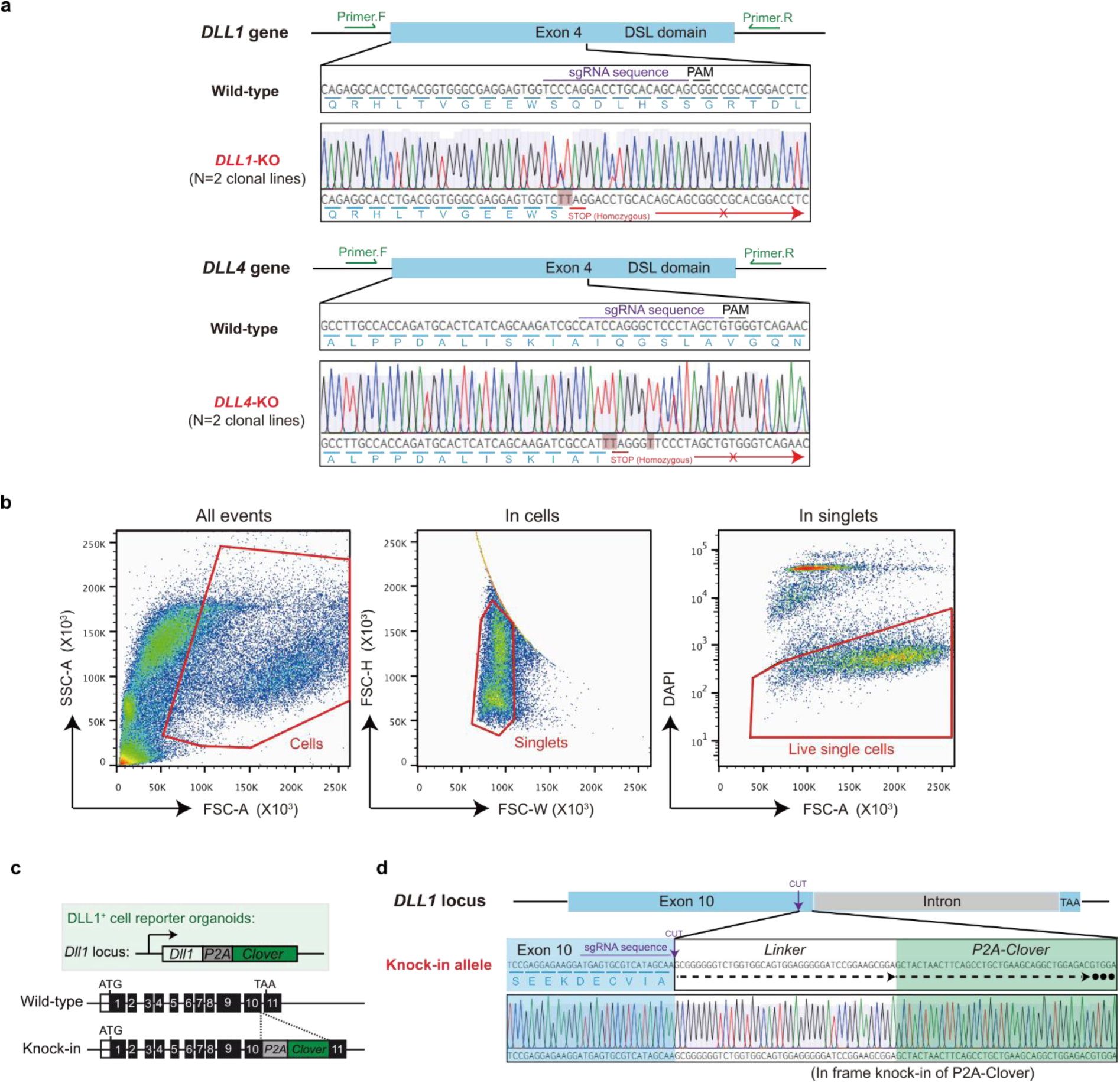
Genotyping and FACS gating strategy of knock-in and knock-out DLL reporter organoid lines. a, Validation of *DLL1* and *DLL4* knock-out (KO) organoids by targeted genotyping. For each targeted gene, n = 2 different homozygous clonal lines are generated. The gene locus, sgRNA targeting sites, locations of genotyping primers, and amino acid sequence (including the CRISPR-generated early STOP codon) are indicated. b, Gating strategy for FACS analysis of organoid cells. All detected events are gated based on their sizes to enrich the cell fraction (left). Doublets are then gated out (middle). Live singlets are gated based on the negative staining of DAPI (right). Additional analysis of cell percentage is based on the fluorescence expression of the knock-in (KI) reporter organoids. c, Illustration of the knock-in reporter organoids containing a *P2A-Clover* cassette inserted at the C-terminus, before the stop codon, of the *DLL1* gene. d, Targeted genotyping of *DLL1-P2A-Clover* reporter organoids.

**Extended Data Fig. 8:**
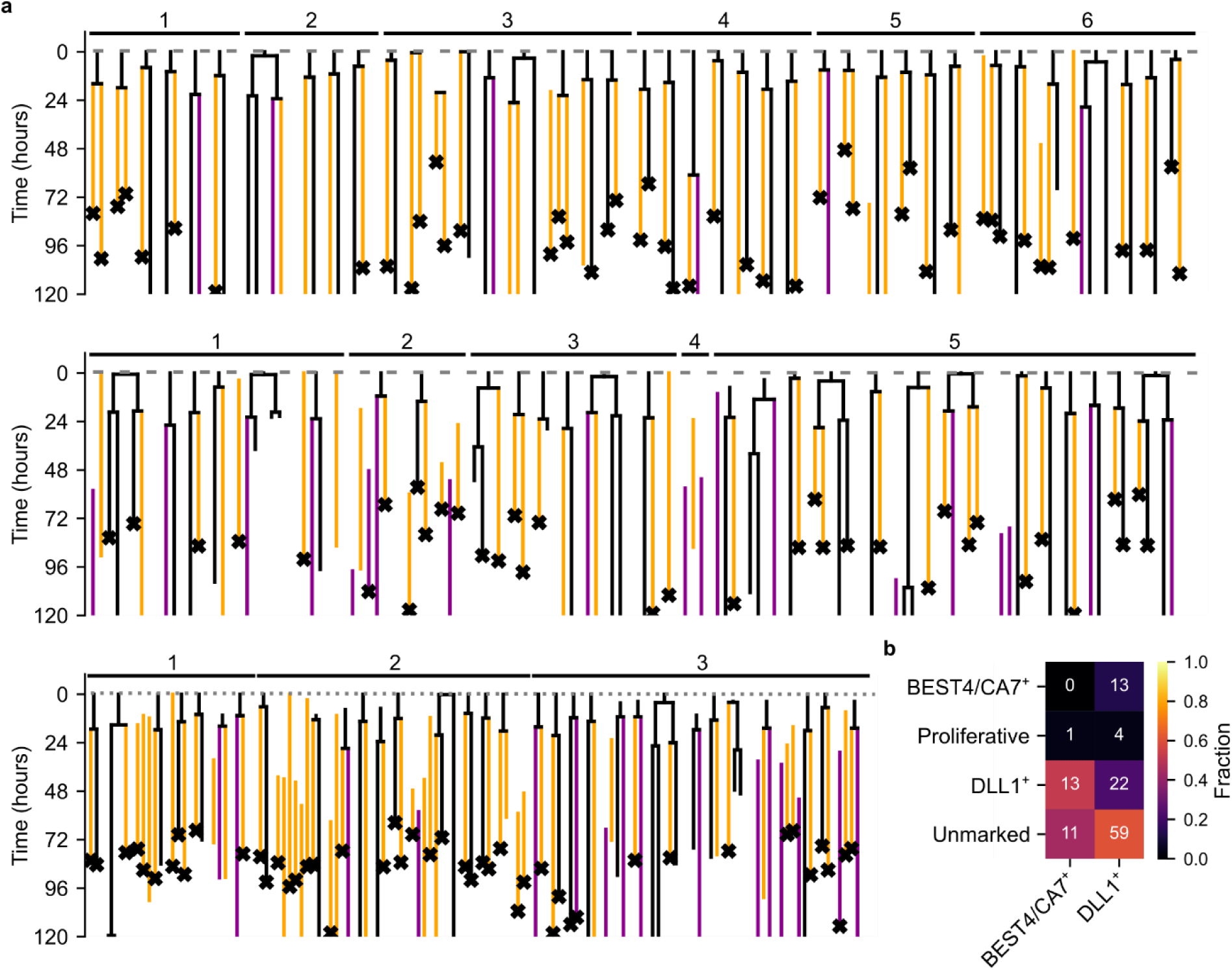
Tracking and lineage tree reconstruction of DLL1^+^ and CA7^+^ cells. a, Lineage tree gallery of CA7-DLL1-H2B reporter organoids. b, Table showing frequencies of sister pairs for the lineages in panel (a).

**Extended Data Fig. 9:**
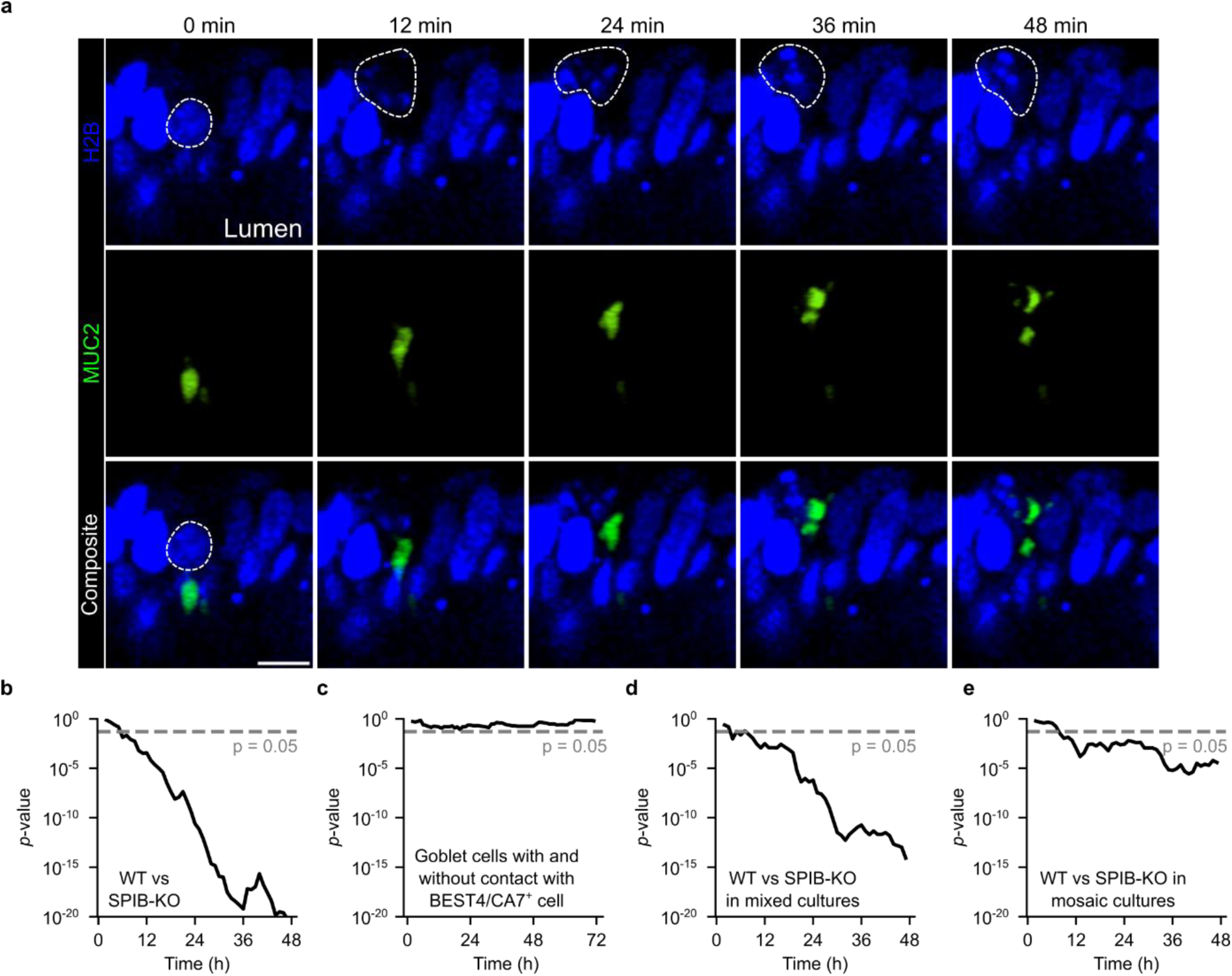
Goblet cell apoptosis. a, Representative example of a goblet cell undergoing apoptosis. Note that the nucleus and the mucus reservoir fragment simultaneously. Scale bar, 10 µm. b-e, Significance (*p*-value) of the comparisons between WT and *SPIB*-KO goblet cells (b), goblet cells with and without contact with a BEST4/CA7^+^ cell at the start of imaging (c), WT and *SPIB*-KO goblet cells in mixed cultures (d), WT and *SPIB*-KO goblet cells in mosaic organoids (e). The *p*-value was computed using the log-rank test with log(− log(⋅)) transformation to increase statistical power^51^.

**Extended Data Fig. 10:**
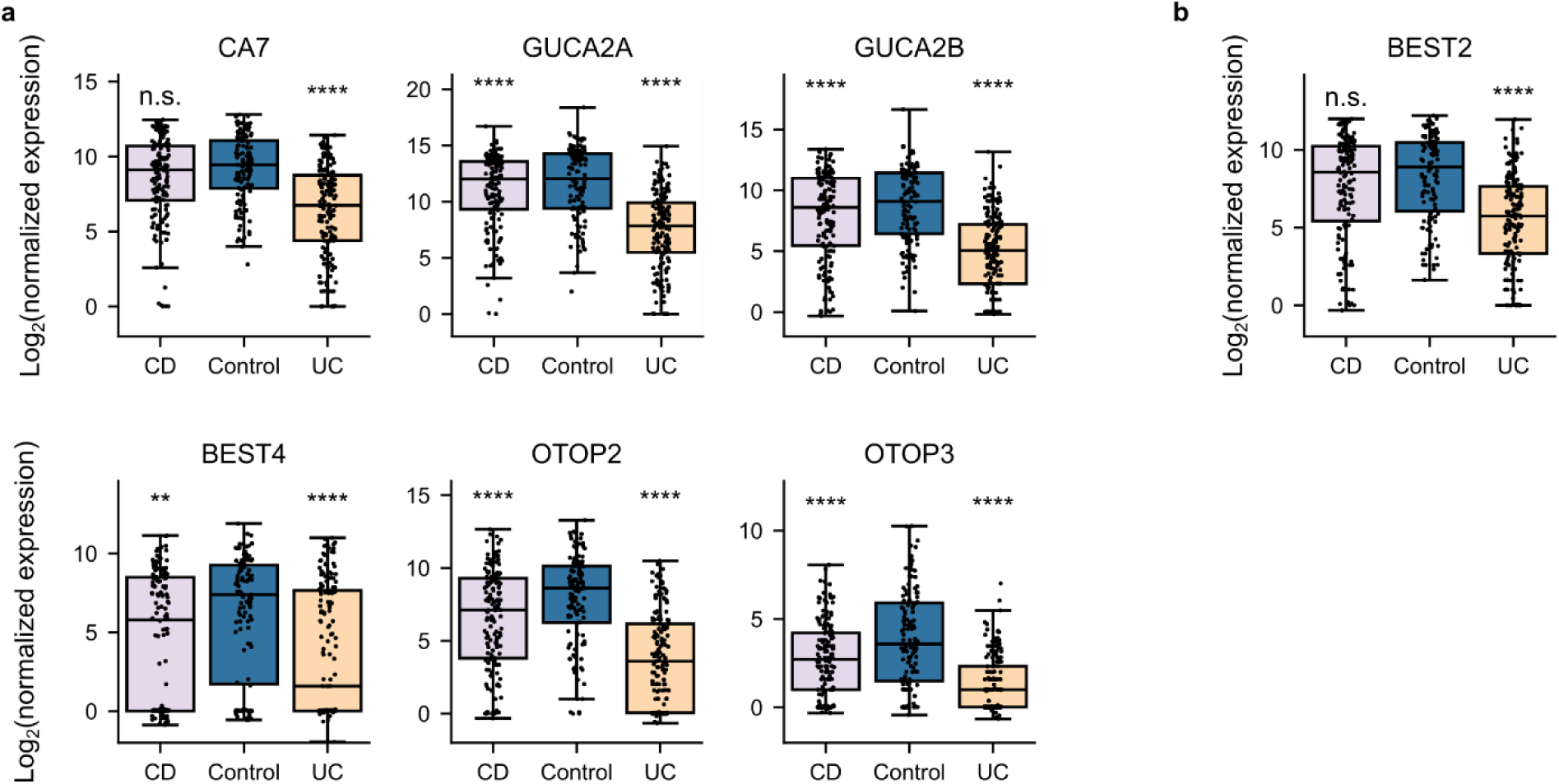
Expression of mature BEST4/CA7^+^ cells and BEST2 is decreased in Ulcerative Colitis and Crohn’s Disease patients. a,b, Expression of mature BEST4/CA7^+^ marker genes (a) and the colonic goblet cell marker BEST2 (b) as retrieved from the IBD TaMMA database^44^. P-values from Tukey’s HSD test were also retrieved from the TaMMA database. **** p < 0.0001, *** p < 0.001, ** p < 0.01, * p < 0.05, and n.s.: not significant. In the box plots, center lines show the median, boxes span the interquartile range (IQR), and whiskers extend to the most extreme data points within 1.5×IQR from the quartiles.

**Supplementary Table 1.**
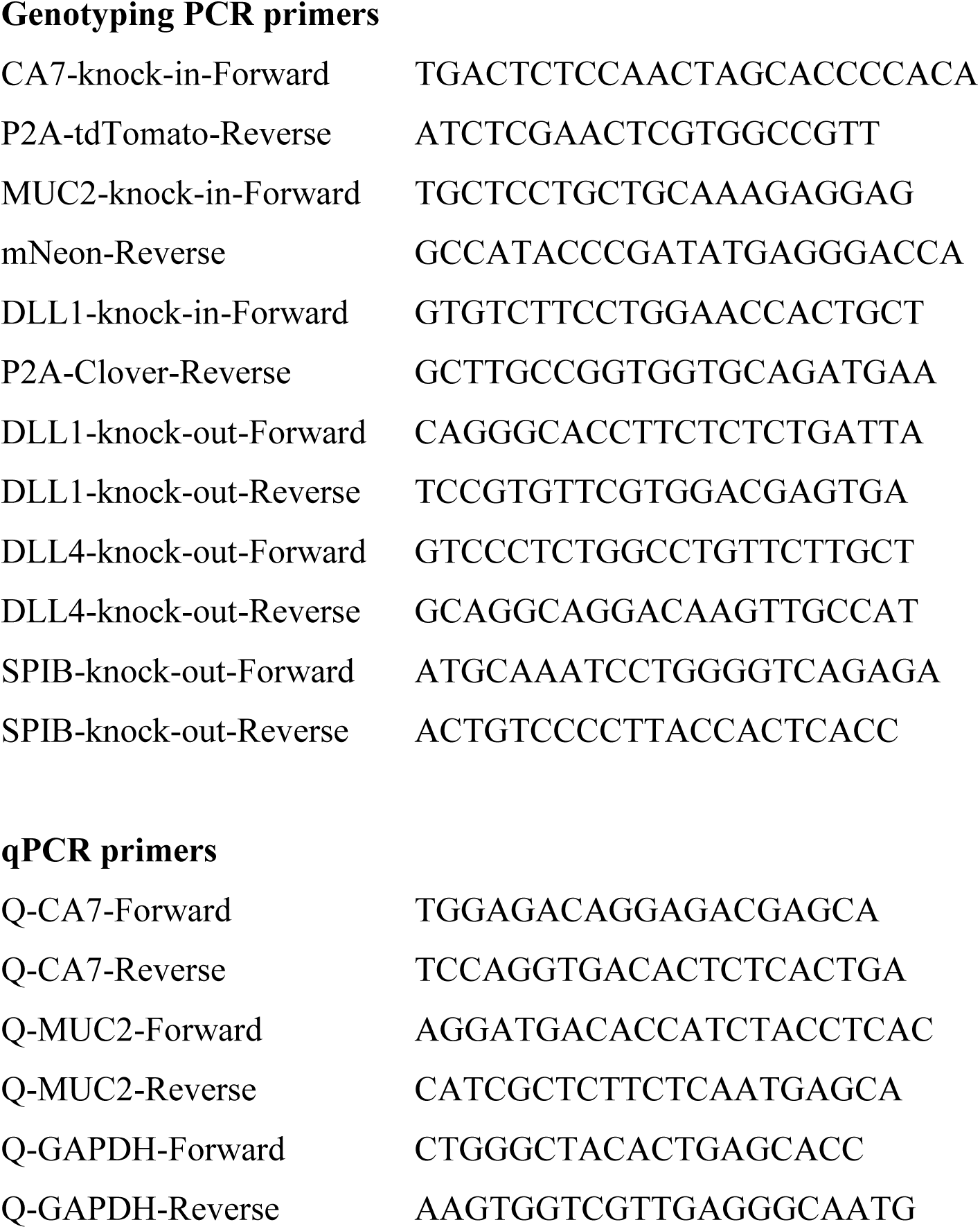
Genotyping and qPCR Primers used in this study.

## Supplementary Videos

Supplementary Video 1: Expansion of reporter organoid. H2B-iRFP670 in grayscale. Scale bar, 20 µm.

Supplementary Video 2: Differentiation of reporter organoid with transcriptional reporters for BEST4/CA7^+^ and goblet cells. MUC2-mNeonGreen reporter in green, CA7- P2A-tdTomato in magenta and H2B-iRFP670 in blue. Scale bar, 20 µm.

Supplementary Video 3: Goblet cell survival in WT organoid. MUC2-mNeonGreen reporter in green, CA7-P2A-tdTomato in magenta and H2B-iRFP670 in blue. Scale bar, 20 µm.

Supplementary Video 4: Goblet cell survival in *SPIB*-KO organoid. MUC2-mNeonGreen reporter in green, CA7-P2A-tdTomato in magenta and H2B-iRFP670 in blue. Scale bar, 20 µm.

